# Colon cancer and cell transformation by clinical *Salmonella* strains are associated with bacterial virulence and intracellular fitness

**DOI:** 10.1101/2023.10.19.562874

**Authors:** Virginie Stévenin, Claudia E. Coipan, Janneke W. Duijster, Daphne M. van Elsland, Linda Voogd, Angela H.A.M. van Hoek, Lucas M. Wijnands, Lennert Jansen, Jimmy J.L.L. Akkermans, Andra Neefjes-Borst, Eelco Franz, Lapo Mughini-Gras, Jacques Neefjes

## Abstract

Non-typhoidal *Salmonella* (NTS) are facultative intracellular pathogens that are associated epidemiologically and experimentally with colon cancer development. Yet, the driving factors of *Salmonella*-induced cell transformation are mostly unknown. We compared 30 (case) NTS clinical strains isolated from patients who were diagnosed with colon cancer >1 year after NTS infection, versus 30 (control) strains from patients who did not develop colon cancer. While we observed diverse cell invasion and transformation efficiencies among the 60 NTS strains, case strains showed higher transformation efficiency than matching control strains. Genomic and transcriptomic analyses showed that transformation efficiency could not be attributed to specific genomic features, but was associated with gene expression, particularly metabolic genes and regulons. Moreover, high-transforming NTS strains display increased capacity to utilize various nutrient sources, including carbohydrates and amino acids, and grow significantly faster intracellularly than low-transforming NTS. Our results link NTS intracellular virulence to cancer promotion.

**Graphical abstract:** 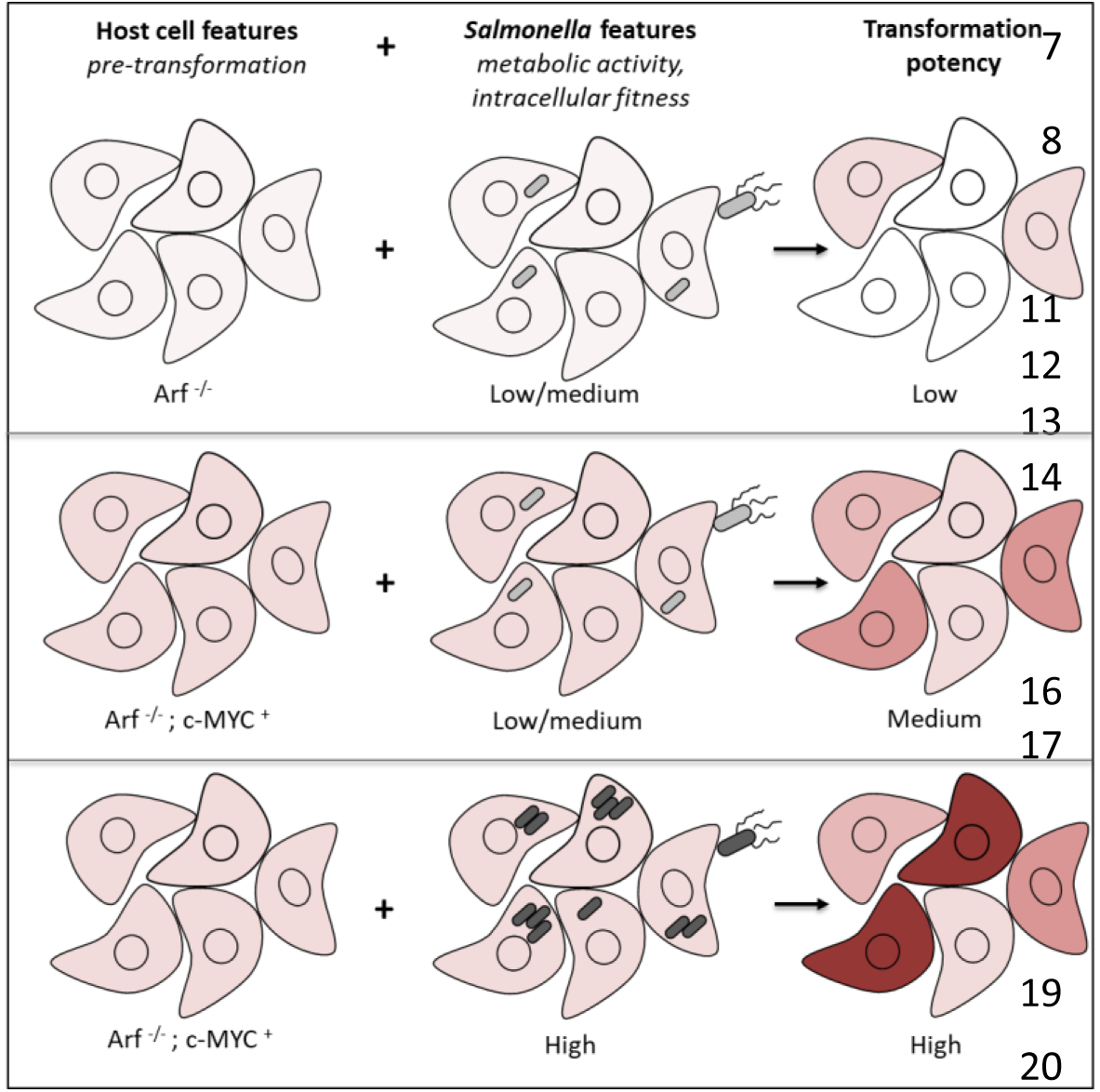

**In brief:** Non-typhoidal *Salmonella* (NTS) infections can promote cell transformation and colon cancer progression. Yet, little is known about the driving factors of *Salmonella*-induced transformation. Stévenin et al. performed a multi-omic characterization of clinical NTS strains identified in a nation-wide epidemiological study as associated with colon cancer and revealed a link between bacterial virulence, intracellular fitness, and host cell transformation.

**Highlights:**

- Cancer-associated clinical NTS generate more cell transformation than matching control NTS.
- NTS transformation efficiency did not correlate with specific genetic features.
- NTS transformation efficiency correlates with gene expression and bacterial metabolic needs.
- High-transforming NTS display increased virulence and intracellular fitness.

## INTRODUCTION

Since the early discovery of *Helicobacter pylori* as a causative agent of gastric cancer (Marshall & Warren, 1984), the role of pathogenic bacteria in the onset and progression of cancers has been gradually acknowledged. While many more bacterial species are added to the list of (potential) oncogenic agents, various mechanisms have been identified by which bacteria directly or indirectly manipulate their host cells promoting cell transformation (van Elsland and Neefjes, 2018). *Salmonella enterica* is a Gram-negative bacteria and a facultative intracellular pathogen. By colonizing a broad range of hosts, including humans, animals, and plants, the circa 2600 *S. enterica* subspecies *enterica* serovars can spread within many niches and are transmitted to humans through contaminated food and water, as well as direct contact with colonized animals, the environment, and via person-to-person transmission. As a result, *Salmonella* is a leading cause of morbidity and mortality worldwide, impairing more than 21 million years of healthy life annually (Kirk et al., 2015).

*Salmonella* serovars are divided between human-restricted typhoid fever-causing serovars (Typhi and Paratyphi), and non-typhoidal *Salmonella* (NTS) serovars. The latter are zoonotic agents with a broader host range, causing mainly gastrointestinal illness in humans (Knodler and Elfenbein, 2019). Following ingestion, a small percentage of salmonellae survive through the gastrointestinal tract, colonize the gut, and invade several host cell types, including the intestinal epithelial cells, where the bacteria can survive and replicate (Galan, 2021). These infections can result in either self-limiting gastroenteritis or invasive diseases. Notably, life-threatening systemic infections associated with typhoid and paratyphoid fever affected 14 million people and led to 130,000 deaths in 2017, mostly in low and middle-income countries (GBD 2017 Typhoid and Paratyphoid Collaborators, 2019). In comparison, NTS serovars cause many more infections worldwide (around 93.8 million), in particular, due to their capacity to spread through the food chain (Majowicz et al., 2010). In most cases, NTS colonization is restricted to the gut and generates mild symptoms allowing recovery without specific treatment (Hohmann et al., 2001). However, life-threatening invasive non-typhoidal salmonellosis is observed in immunocompromised individuals, in particular in sub-Saharan Africa and Southeast Asia (Stanaway et al., 2019).

In the last decade, evidence provided by us and others support that NTS infections increase the risk of colon cancer development (Scanu et al., 2015; Mughini-Gras et al., 2018; Zha et al., 2019; van Elsland et al., 2022). To thrive within its host cell, *Salmonella* hijacks many intracellular signaling pathways (Galan, 2021), some of which may result in host cell transformation (Scanu et al., 2015; Lu et al., 2016). We have previously shown that *Salmonella* Typhimurium invasion of host cells can induce cell transformation in tissue culture (Scanu et al., 2015). However, it is unclear whether NTS strains differ in their transformation efficiency.

In an earlier epidemiological study, we have shown that colon cancer risk was significantly increased among patients with a severe – therefore reported - NTS infection diagnosed before 60 years of age (Mughini-Gras et al., 2018). In particular, infection with NTS serovar Enteritidis was associated with the highest increased risk of cancer in the ascending/transverse colon (a 3-fold increase compared to no infection), suggesting different transformation efficiency between NTS strains (Mughini-Gras et al., 2018). These epidemiological insights were the basis for the current study. Here, we show that tumors linked to NTS infections have a different pathological grading than colon tumors from patients without reported NTS infection. We further describe the NTS features associated with colon cancer performing a matched case-control study of clinical NTS strains either or not associated with colon cancer as identified in the nationwide epidemiological study (Mughini-Gras et al., 2018). Using a combination of genomic, transcriptomic, metabolic, and phenotypic analyses of the clinical NTS strains, we report a surprisingly diverse efficiency of clinical NTS to transform host cell and describe how this attribute correlate with strain transcriptome, virulence, and intracellular fitness.

## RESULTS

### Tumors from patients with a history of reported *Salmonella* infection are associated with a better prognosis

Since 1991, a nationwide histopathology and cytopathology network and archive has been in operation in The Netherlands (PALGA), encompassing all sixty-four Dutch pathology laboratories (Casparie et al., 2007). We coupled the data from our previous epidemiological study (Mughini-Gras et al., 2018) to that of PALGA to retrieve tissue blocks corresponding to the patients with colon cancer that were registered at the Dutch National Institute for Public Health and the Environment (RIVM) with a diagnosed NTS infection earlier in life (SL^+^) or not (SL^-^). Both groups were matched in age, gender, and tumor localization (proximal versus distal colon). Using PALGA registration, we localized and retrieved the corresponding tissue blocks through the network of pathology laboratories. We obtained 24 SL^+^ and 67 SL^-^ colon tumor blocks. The quality of the tissue blocks varied strongly, allowing only Hematoxylin and Eosin (HE) staining and evaluation by an experienced pathologist at the Amsterdam University Medical Center (Amsterdam UMC). The evaluation was performed in a blinded manner. Tumor grading showed a tendency for the SL^+^ tumors to be more differentiated – corresponding to a better prognosis - than SL^-^ tumors (odds ratio 0.21, 95% CI 0.004-1.06; p-value 0.059; Figure 1). These differences did not reach full statistical significance, likely due to the expected presence of background noise in both SL^+^ and SL^-^ groups. Indeed, the epidemiological definitions of the groups SL^+^ and SL^-^ are correlative but not causal, meaning that the SL^+^ tumor also includes colon cancer development independent of NTS infection. Besides, the patients with SL^-^ tumors may have been exposed to NTS while developing no or mild symptoms. Those most commonly mild and unreported infections are sufficient to promote colon cancer (van Elsland et al., 2022) but are not registered as patients with an NTS infection. Despite these intrinsic limits, the outcomes of our pathology analyses are aligned with earlier research showing that colon tumors of patients with a history of NTS infection were mostly of low grade (Mughini-Gras et al., 2018). These results suggest a different biology between tumors associated or not with NTS infection.

**Figure 1:**
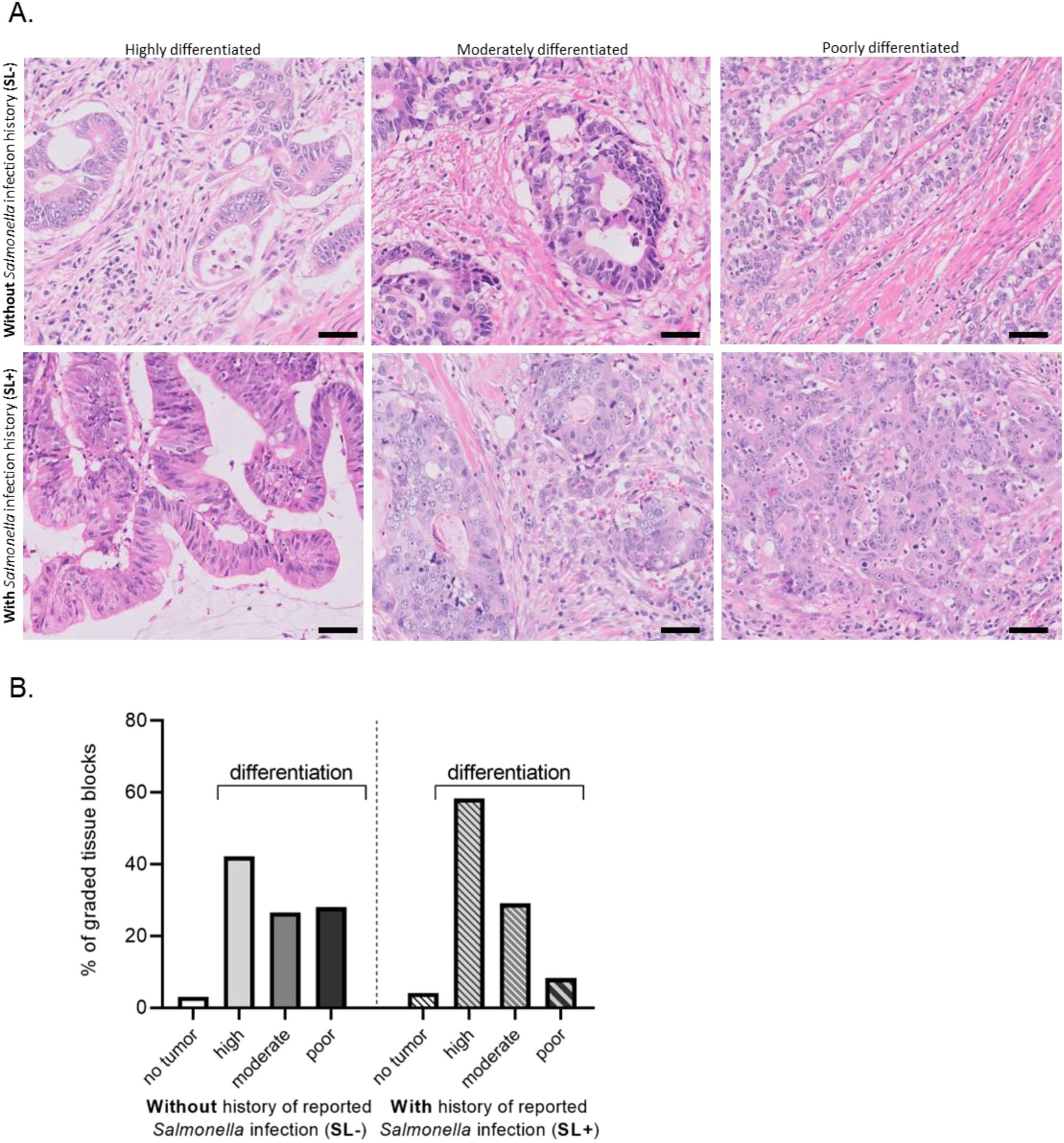
Tumor staging of colon tumor blocks from patients with and without a history of reported NTS infection. **A.** Representative images of the three levels of tumor grading (highly, moderately, and poorly differentiated) of the colon tumor blocks from patients with (SL^+^) and without (SL^-^) a history of reported NTS infection. Sample stained with hematoxylin and eosin. Scale bar: 50 µm. **B.** Distribution of tumor staging (in percentage) of the colon tumor blocks from patients with (SL^+^, n=24) and without (SL^-^, n=67) reported history of NTS infection. Odds ratio: 0.21; 95% CI: 0.04-1.06; p-value: 0.059. *- back to text.*

### Transformation efficiency varies between clinical NTS strains

We aimed to identify potential features of clinical NTS strains that could promote tumor development. We retrieved clinical NTS strains associated with colon cancer utilizing previously linked data from the Dutch national surveillance system for *Salmonella* and the Netherlands Cancer Registry (Mughini-Gras et al., 2018). Clinical NTS strains were sent from peripheral medical microbiological laboratories to the RIVM for further typing and storage (Van Pelt, 2003). Using this collection, we defined as “case strains” (Figure 2A, red salmonellae) 30 clinical NTS strains isolated from patients with a reported NTS infection who developed proximal colon cancer at least one year after the reported salmonellosis in the period 2000-2015. In addition, we defined as “control strains” (Figure 2A, green salmonellae) 30 clinical NTS strains from the same *Salmonella* surveillance database, which were obtained from patients who did not develop colon cancer after infection (until at least the end of 2015). The control strains were selected to match (1:1 ratio) the case strains regarding serovar, type of infection (enteric, septicemic, etc.), year of infection, age at infection, and gender of the patient (Figure 2A-B, Supplementary Table S1).

**Figure 2:**
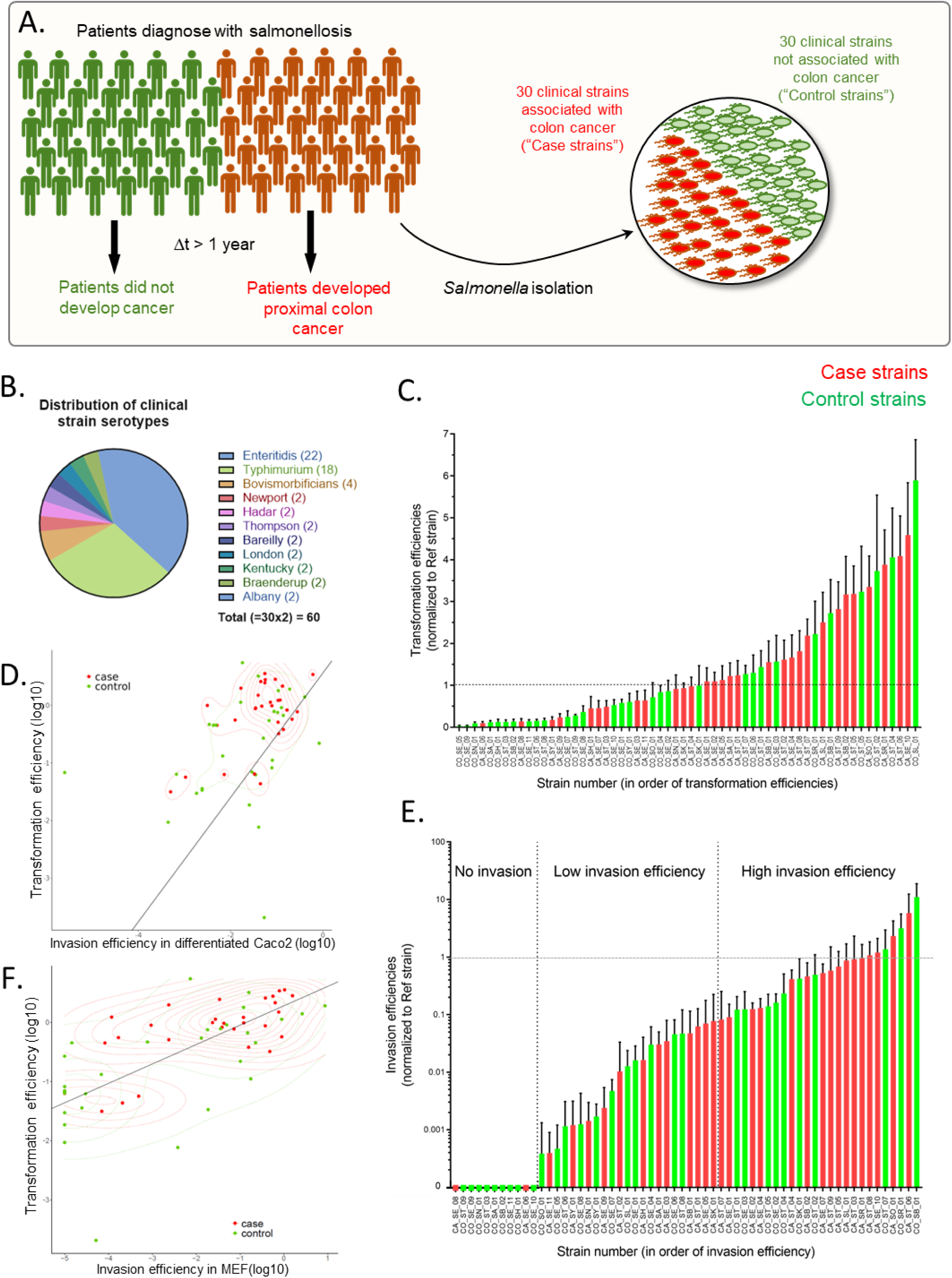
Clinical NTS strains related to cancer development and control patients differ in both transformation and invasion efficiencies. A. Schematic of the clinical strain selection. 30 clinical strains isolated from patients who developed colon cancer >1 year post-infection (“case strains”, in red) were matched with 30 clinical strains isolated from patients without cancer development (“control strains”, in green). B. Serovar distribution of the 60 clinical NTS strains. C. Strain transformation efficiency in MEF assessed by soft-agar assay, normalized by the transformation efficiency of the reference strain, and ordered by transformation efficiency. Error-bar: SEM. D. Strain transformation efficiency in MEF (mean) as a function of invasion efficiency in Caco2 cells (mean). Grey line: orthogonal regression. Green and red dashed lines: density of control and case values, respectively. E. Strain invasion efficiency in MEF assessed by gentamicin assay, normalized by the infection efficiency of the reference strain, and ordered by infection efficiency. Error-bar: SEM. F. Strain transformation efficiency (mean) as a function of invasion efficiency in MEF (mean) as measured in C and E. Grey line: orthogonal regression. Green and red dashed lines: density of control and case values, respectively. *- back to text.*

We previously reported that *Salmonella*-induced cell transformation required at least two host pre-transforming steps before infection, in particular the inactivation of the p53 pathway (by knocking out Arf) and the overexpression of the proto-oncogene c-MYC (Scanu et al., 2015). We thus performed an *in cellulo* characterization of the 60 clinical strains in mouse embryonic fibroblasts (MEF) overexpressing c-MYC and *Arf*-deficient, a model cell line that allows monitoring transformation with classical anchorage-independent colony-forming assays (Scanu et al., 2015; van Elsland et al., 2022; Stévenin and Neefjes, 2023). We normalized the results to the transformation efficiency of the lab strain *S*. Typhimurium SL1344 (later referred to as “reference strain”). We observed that 18 case strains (representing 60% of all case strains) and 10 control strains (representing 33% of all control strains) showed a higher transformation efficiency than the reference strain (Figure 2C, Supplementary Table S2). On average, the case strains had a higher transformation efficiency compared to the control strains (1.66 vs. 1.15). The ratio between the paired values of the matching case-control strains was significantly different from 1.0 (ratio paired t-test, p=0.0123), meaning that case strains had a significantly higher transformation efficiency than their control strains (Figure S1A). Thus, our results suggest that high transformation efficiency is a broadly spread feature of clinical NTS strains, also reflected by their clinical association with colon cancer.

### Transformation efficiency correlates with invasion efficiency

We assessed the intrinsic capacity of the 60 clinical NTS strains to reach and invade their host cell. First, we used a 2-step *in vitro* gastrointestinal tract (GIT)-model to test the intrinsic capacity of the strains to survive during their passage through the GIT. For this, overnight cultures (ON) of NTS were diluted and incubated in a simulated gastric fluid (SGF) at 37 °C for 30 min, mimicking the gastric passage. Then, SGF cultures were diluted and incubated in a simulated intestinal fluid (SIF) at 37 °C for 2 h, mimicking the intestinal environment (Kuijpers et al., 2019; Figure S1B). We scored the SGF/ON, SIF/ON (Figure S1C-D) and SIF/SGF OD ratios for all strains (Supplementary Table S3). These ratios appeared independent from the strains’ *in cellulo t*ransformation efficiency based on a linear regression analysis suggesting that the GIT survival and transformation efficiency are intrinsic independent strain features. Besides, no statistically significant difference was observed between case and control strains based on conditional logistic regression analysis (Supplementary Table S3). We then tested the capacity of the strains to attach to (ATT) and invade (INV) polarized enterocytes by performing gentamicin protection assays after infection of differentiated epithelial Caco2 cells. While there was again no significant difference between the case strains and control strains based on conditional logistic regression analysis, we found that INV/ON and INV/ATT ratios are correlated with the transformation efficiency (correlation coefficients: 0.41 and 0.32 respectively; orthogonal regression in Figure 2D; Supplementary Table S3).

In addition, we tested the invasion efficiency of the 60 clinical strains in the MEF model (Figure 2E, Supplementary Table S2), normalized by the invasion efficiency of the reference strain. Among the 60 clinical NTS strains, 24 had an invasion efficiency equivalent to or higher than the reference strain, 25 had an invasion efficiency at least 10 times lower than the reference strain, and 11 did not infect the MEFs (Figure 2E). While the invasion efficiency of the control and case strains were not significantly different (t-test p>0.05), the strains that could not invade the MEF mostly belonged to the control group (82%, Figure 2E, Supplementary Table S2). Additionally, the transformation efficiency was significantly correlated to the invasion efficiency in MEFs (orthogonal regression in Figure 2F; correlation coefficient: 0.67). These results are consistent with our previous observation that cell invasion is required for *Salmonella*-induced cell transformation (Scanu et al., 2015), and further link transformation efficiency and virulence.

### Investigation of genomic features associated with transformation efficiency

To determine whether there were genetic markers that could explain the variability in transformation efficiency of the *Salmonella* strains, we performed several genome-wide association tests (GWAS) using all or a subset of the genomes of the 60 different clinical NTS. We observed that the presence or absence of plasmids (Figure 3A) among the 60 NTS strains did not correlate with NTS transformation efficiency. We further analyzed the presence/absence of 233 *Salmonella* virulence genes. 128 virulence genes were present in all NTS strains investigated, whereas 26 were found in none (Figure S2A). This latter result concerned operons typical for *S*. Agona and *S*. Typhi, serotypes not included in the present study. Similarly to plasmids, variations in the presence/absence of virulence genes between the 60 strains did not correlate with NTS transformation efficiency (Figure S2A).

**Figure 3:**
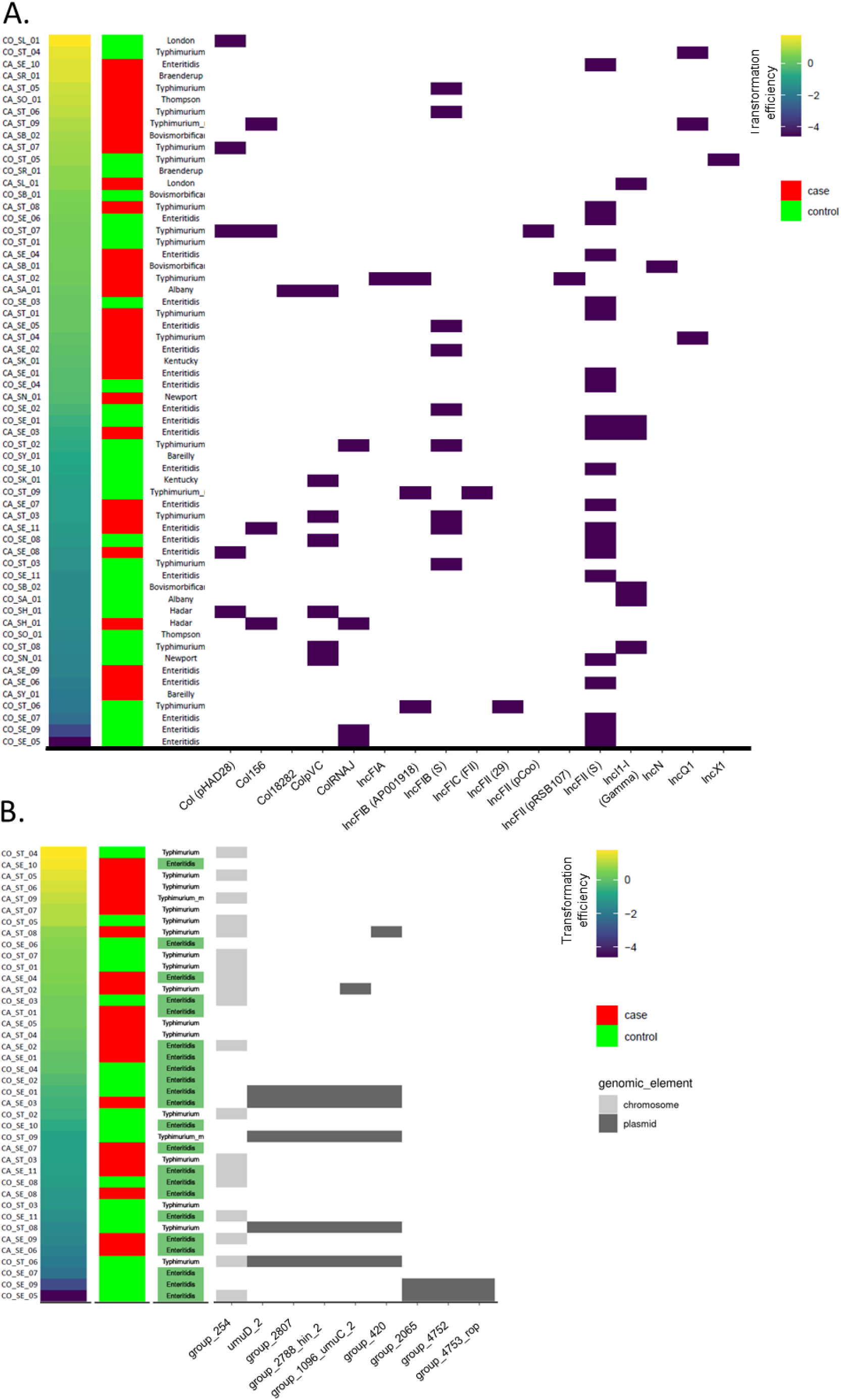
NTS strain transformation efficiency does not correlate with specific genomic features. A. The presence of plasmids (x-axis, purple boxes) among all 60 strains in the study did not correlate with the strain transformation efficiency (y-axis). The first three columns in the panels correspond to the transformation efficiency (log-median values), type of strain (control or case), and bacterial serovar. B. Distribution of the genes identified as significantly associated with the transformation efficiency by TreeWAS, among the *S*. Typhimurium and *S*. Enteritidis strains. The first four columns in the panels correspond to the transformation efficiency (log-median values), type of strain (control or case), bacterial serovar (*S*. Typhimurium or *S*. Enteritidis) and location of genomic elements (chromosome or plasmid). The gene names are based on the output of the pan genome pipeline Roary. *- back to text.*

For more in-depth genetic comparison, we restricted the analysis to the combination of the two most abundant serovars: *S*. Enteritidis and *S*. Typhimurium, corresponding to 22 and 18 clinical strains respectively (while other serovars had only 2-4 clinical strains each, limiting statistical power). Five genes appeared significantly associated with higher transformation efficiency in the *S*. Typhimurium subset and four in the combined *S*. Enteritidis/Typhimurium subset (Table 1, Figure 3B). One of the significant genes of the *S*. Typhimurium subset and three from the combined *S*. Enteritidis/Typhimurium subset had unknown functions (Table 1, Figure 3B). For the remaining genes, functional annotation revealed three proteins involved in UV protection and mutation of which two are part of the bacterial SOS response to DNA damage (UmuC and UmuD) (Table 1, Figure 3B). Eight out of nine of these genes are localized on plasmids. While the presence/absence of several genomic elements statistically correlates with clinical strain transformation efficiency, these elements were broadly present within high and low transformers, or only present in low transformers (Table 1, Figure 3B), making their implication in bacterial-induced transformation doubtful.

**Table 1.** Genes significantly associated with transformation efficiency in *S*. Enteritidis, *S*. Typhimurium, and combined *S*. Typhimurium/*S*. Enteritidis subsets. back to text.

| Gene name | Annotation | Gene function |
| --- | --- | --- |
| ***S. Typhimurium* subset** |  |  |
| group_2788 | DNA-invertase hin<br>[ https://www.uniprot.org/uniprot/P03013 ] | Synthesis of phase-2 flagellin. |
| group_2807 | protein ImpC<br>[ https://www.uniprot.org/uniprot/P0A1G0 ] | Belongs to the imp operon which has a function in UV protection and mutation. |
| group_1096 | protein UmuC | Functions in UV protection and mutation, induced/SOS mutation. |
| group_420 | *unknown* | *unknown* |
| umuD_2 | protein umuD<br>[ https://www.uniprot.org/uniprot/P22493 ] | Functions in UV protection and mutation, induced/SOS mutation. |
| **Combined *S. Typhimurium* and *S. Enteritidis* subset** |  |  |
| group_2065 | *unknown* | *unknown* |
| group_2540 | *unknown* | *unknown* |
| group_4752 | *unknown* | *unknown* |
| group_4753 | regulatory protein rop<br>[ https://www.uniprot.org/uniprot/P03051 ] | Regulatory role in plasmid DNA replication. |

Additionally, we compared the genome organization of the *Salmonella* pathogenicity islands (SPI) 1, 2, and 4 encoding the type 1 and 3 secretion systems (T1SS and T3SS) within 3 *S*. Typhimurium and 3 *S*. Enteritidis clinical strains. We observed similar - if not almost identical - structures for both *S*. Typhimurium and *S*. Enteritidis (Figure S2B). Lower identity scores observed in SPI4, in an open reading frame (ORF) corresponding to an unknown protein, were due to low complexity genic regions that allow multiple matches between sequences. However, the protein sequences were heavily conserved for the majority of the clinical strains. Thus, we could not identify a convincing genetic feature associated with the transformation efficiency of clinical NTS strains.

### Transcriptomic analyses link NTS transformation efficiency to high expression of particular metabolic genes

The lack of genetic features correlating with transformation efficiency raised the question of whether different gene expression levels could be associated with transformation efficiency. During its intracellular life, *Salmonella* regulates the transcription of specific genes necessary for the bacteria to enter, survive, and replicate in the eukaryotic cells (Srikumar et al., 2015; Powers et al., 2021). To characterize the transcriptomic profile of NTS in association with their transformation efficiency, we selected eight NTS strains, corresponding to two high- and two low-transforming strains from both *S*. Typhimurium and *S*. Enteritidis with maximal genetic proximity (Figure S4A). The strains were grown under *in vitro* conditions promoting the induction of SPI1 or SPI2, as previously demonstrated (Kröger et al., 2013; Lober et al., 2006; Srikumar et al., 2015). The SPI1-inducing growth condition expectedly enables the induction of the *Salmonella* transcriptomic program as observed at the invasion and early stage of the infection (Kröger et al., 2013). Once intracellular, *Salmonella* typically resides in a low pH *Salmonella*-containing vacuole (SCV) triggering SPI2 expression. While most SCV are turned into vacuolar replicative niches, about 10-20% of the SCV ruptures, leading to *Salmonella* release in the host cytosol (Gutierrez & Enninga, 2022). The SPI2-inducing growth condition mimics the intravacuolar acidic low-phosphate environment and thus induces the *Salmonella* transcriptomic program as observed in the SCV from 1-2 h post-invasion (Löber et al., 2006). In addition, as similarities in gene expression in extracellular and cytosolic bacteria, in particular SPI1, have been reported, we suspect that the SPI1-inducing condition also partially summarizes cytosolic bacteria gene expression (Powers et al., 2021).

Following RNA extraction and sequencing, the number of reads obtained varied extensively among and within the bacterial strains and experimental conditions, with the median number of reads mapping to coding DNA sequences (CDS) ranging between 6.2 and 7.8 million in the SPI1-inducing growth condition and between 9.2 and 17.6 million in the SPI2-inducing growth condition (Figure S3). We confirmed that the *in vitro* growth in SPI1- or SPI2-inducing conditions indeed led to the expression of SPI1 and SPI2, respectively, in the studied strains (Figure S4B-C). We then analyzed the differential gene expression between the strains with high and low transformation efficiencies and the corresponding pathway enrichment in each of the two experimental conditions. There was a low number of genes differentially expressed in the SPI1-inducing growth condition (n = 14 genes, Supplementary Table S4), and a significantly higher number in the SPI2-inducing growth condition (n = 387, Supplementary Table S4). This might indicate that the low- and high-transforming strains behave similarly under the SPI1-inducing condition and only show a different behavior under the SPI2-inducing condition. A vast majority of the genes overexpressed in the SPI2-inducing condition (n = 184) showed more than a log2-fold change (i.e. doubling; Supplementary Table S4). About half of the over-expressed genes (n = 201/401) were associated with a KEGG metabolic pathway (Figure 4A-B; Supplementary Table S5). KEGG pathway enrichment analyses (Supplementary Table S6) indicated that numerous metabolism-associated genes were over-expressed in the SPI2-inducing growth condition, with carbohydrate, amino acid, and cofactors and vitamin metabolism highly active. Additionally, transport across membrane and cell motility pathway classes were enriched (Figure 4A-B). This suggests that a higher metabolic activity accompanies the intravacuolar phase of the bacterial infection. This finding was consistent with a previous report of increased expression of metabolic genes by *Salmonella* in macrophages (Srikumar et al., 2015). Of the differentially expressed genes upon SPI2 induction 148 were grouped in 50 operons, and 94 of these (grouped in 29 operons) were over-expressed in the bacterial strains with a higher transformation efficiency; the dominant operons were involved in flagellar assembly (*flg* and *fli* genes) and chemotaxis (*che*, *mot*, and *tar* genes; cell motility, signal transduction), thiamine metabolism (*thi* genes; metabolism of cofactors and vitamins), pilus assembly (*yeh/stc* genes), oxidative phosphorylation (*cyo*, *nuo* and *ndh* genes; energy metabolism), citrate cycle (*sdh* genes; TCA cycle, carbohydrate metabolism), and ABC transporters (*met/sfb* genes; membrane transport). Among the genes that were over-expressed in the bacterial strains with low transformation efficiency were the *kdp* genes (involved in potassium transport), the *lsr* genes (ABC transporters), and the *thr* genes (involved in amino acid metabolism and biosynthesis) (Supplementary Tables S4, S5, S6). Overall, we observed cell motility, energy metabolism, metabolism of cofactors and vitamins, and signal transduction pathways to be particularly enriched in high transformers while quorum sensing genes were overexpressed in low transformers.

**Figure 4:**
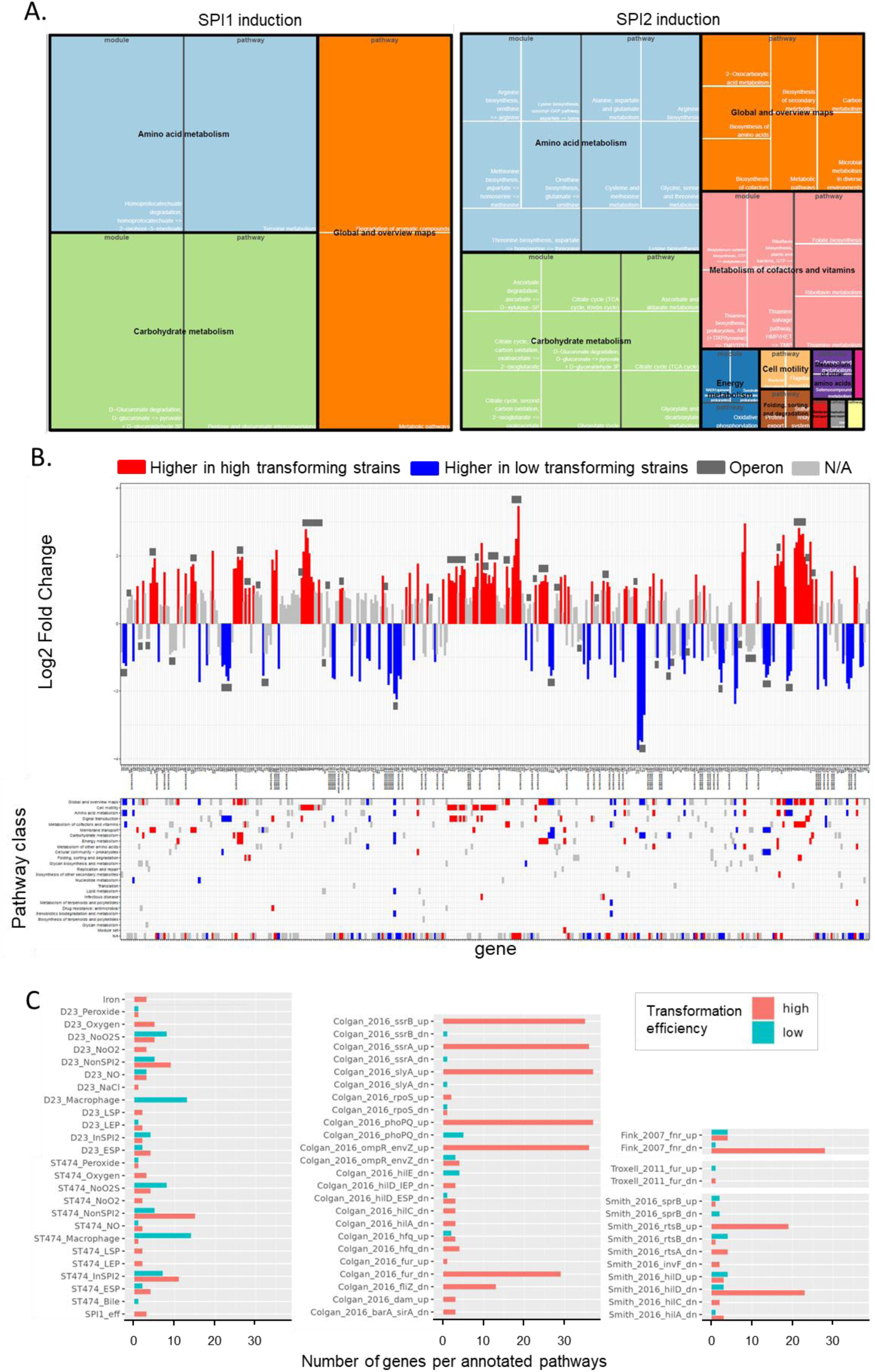
NTS transformation efficiency correlates with specific transcriptomic profile. A. KEGG module and pathway enrichment upon SPI1 (left tree map) and SPI2 (right tree map) inductions in both *S*. Enteritidis and *S*. Typhimurium. The size of the tiles represents the fraction of overexpressed genes identified in each module and pathway. Links to high-resolution tree maps: [left](https://drive.google.com/file/d/1DXItyAPQ042_MooO9IX9_9Mq97z4x74G/view?usp=sharing), [right](https://drive.google.com/file/d/1coKoj72LZf-iNpOno5ATUnXFuBYitmk3/view?usp=sharing). B. Increase (log2 fold change) in gene expression (top panel) and associated pathways (bottom panel). The vertical bars in the upper panel indicate the magnitude of change in gene expression of one category compared to the other (I.e. high and low-transforming strains), with the genes significantly overexpressed in high transformers above the x-axis and the ones significantly overexpressed in low transformers under it. The color of the vertical bars in the upper panel and horizontal ones in the lower panel indicates genes with a log2 fold change > 1 in high transformers (red), genes with a log2 fold change < 1 in low transformers (blue), and genes with a log2 fold change < 1 in any of the high of low transformers (light grey). Horizontal dark grey bars in the upper panel indicate operons. [high-resolution panel](https://drive.google.com/file/d/1PnlKU9my7HqGg9dcwLOD3N6ZVhF53XEM/view?usp=sharing). C. Pathway enrichment analysis within regulons generated from transcriptional profiling of key regulatory mutants (up: upregulated; dn: downregulated) or from RNA-seq profiling of *S*. Typhimurium under 22 infection-relevant *in vitro* growth conditions (annotated by Powers et al., 2021), in bacteria with high (red) and low (blue) transformation efficiency grown in SPI2-inducing condition. *- back to text.*

To further define the transcriptomic profile of high- and low-transforming strains, we used the custom annotation by Powers and colleagues of *Salmonella* regulons and RNA-seq profiling observed in different infection-relevant *in vitro* growth conditions (Powers et al., 2021). About a quarter of the over-expressed genes (n = 98/401) were associated with one of these custom pathways (Supplementary Table S5). Pathway enrichment analyses (Supplementary Table S6) indicated that the strains with high transformation efficiency expressed more genes associated with regulons controlled by rtsB, ssrA, ssrB, slyA, phoPQ, envZ (upregulated in mutated strains) and hilD, fnr, fur and fliZ (downregulated in mutated strains, Figure 4C). This divergence in the transcriptomic program of high- and low-transforming bacteria suggests a different response to the environmental cue present in the SPI2-induction growth medium.

### Metabolic profiling using phenotype microarray technology reveals an association between high transformation efficiency and specific metabolic behavior

As the transcriptomic analysis revealed a link between the expression of genes involved in various metabolic pathways and transformation efficiency, we also investigated the metabolic pathways potentially used by the 60 clinical NTS strains with a phenotype microarray (Shea et al., 2012). We tested the capacity of each clinical NTS strain to grow in media that contains one of the 284 tested chemical compounds providing carbon, nitrogen, phosphorus, and sulfur sources to bacteria (Supplementary Table S7, Figure S5). We revealed 76 positive and 8 negative correlations between substrate utilization and transformation efficiency (Supplementary Table S7). We generated a heatmap of the scaled utilization scores of these correlations using average linkage clustering (Figure 5A). The results highlight a tendency towards increased substrate utilization for strains with a higher transformation efficiency, particularly pronounced for amino acids and derivatives such as peptides (Figure 5A; Supplementary Table S7). Of note, some of these substrates correlated with transformation efficiency when used as carbon and nitrogen sources. We compared the outcome of the transcriptomic analyses with the utilization of metabolic substrates, by visualizing the intersection of the pathways enriched in both analyses (Figure 5B). We identified eight pathways that were enriched in both analyses; four of them were associated with amino acid metabolism, three with carbohydrate metabolism, and one with energy metabolism (Figure 5B, Supplementary Table S6).

**Figure 5:**
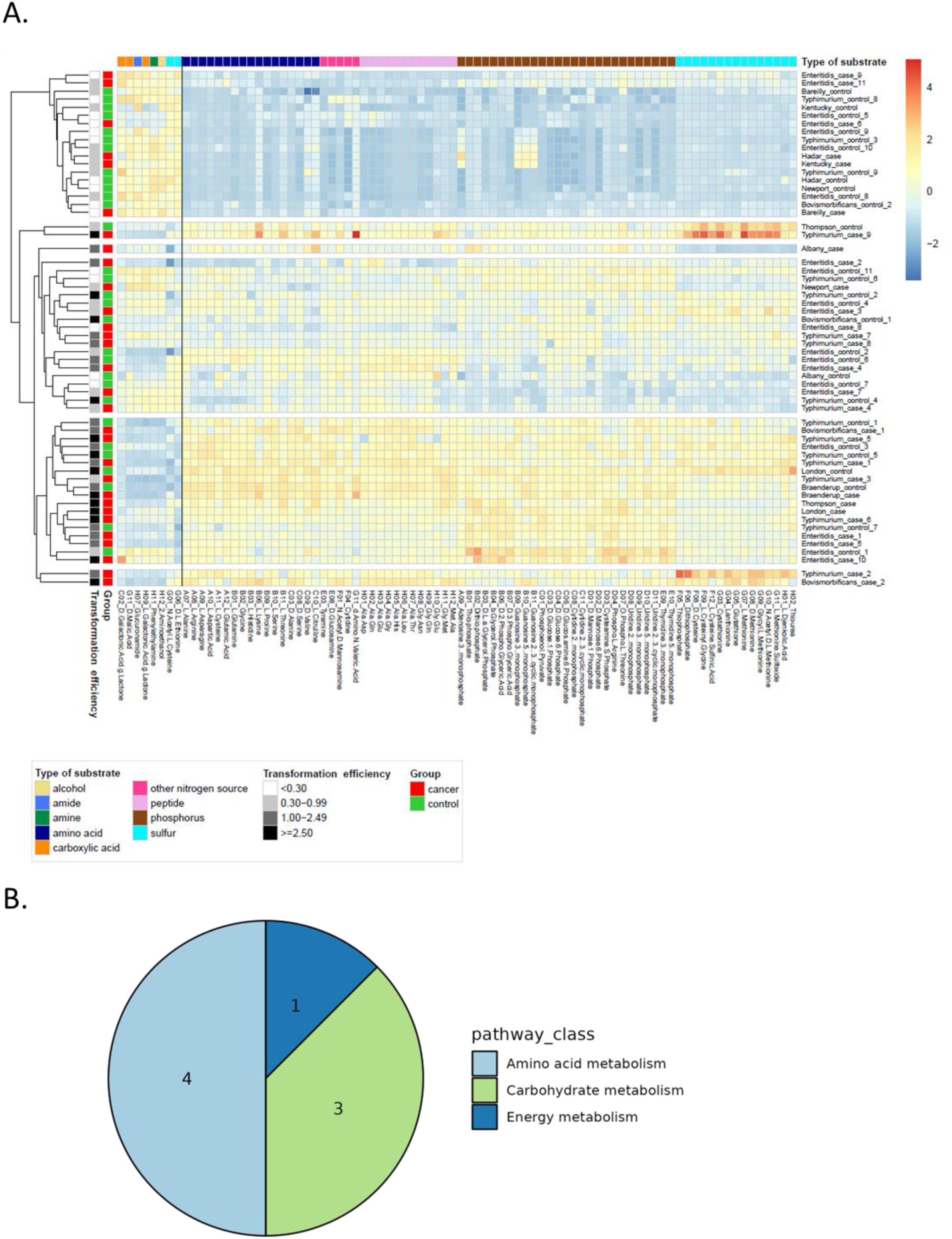
High NTS transformation efficiency correlates with high capacity to consume amino acids and derivatives for growth. A. Heatmap of scaled utilization scores of the 60 NTS clinical strains for the sources with a significant positive or inverse correlation with transformation efficiency. Left of the black line: 8 columns depict the sources with negative correlation, right of the black line 76 columns depict the source with a positive correlation. NTS strains are clustered based on their utilization scores using the average linkage method. B. Pathway classes (KEGG annotation) over-represented in both metabolic and RNAseq analyses *- back to text.*

### High transformation efficiency is associated with high rates of intracellular replication

Given the high amount of amino acid in the intracellular environment, we compared the intracellular behavior and fitness of low and high transformers by confocal time-lapse microscopy. Typically, the nascent SCV undergoes sequential maturation until the establishment of a mature and permissive vacuolar niche from which LAMP1-positive *Salmonella*-induced filaments (SIFs) emanate (Fang & Méresse, 2022). Alternatively, SCV’s rupture can lead to *Salmonella* replicating at a high pace (so-called “hyper-replication”) in the cytosol (Gutierrez & Enninga, 2022). As the capacity of *Salmonella* to hyper-replicate in the cytosol is restricted to certain cell types such as epithelial cells, *Salmonella* has been described as an “opportunistic” cytosolic pathogen (Powers et al., 2021). Importantly, the nutrient sources obtained by *Salmonella* depend on its vacuolar or cytosolic localization, as well as the formation of SIFs (Kehl et al., 2020).

To characterize the intracellular behavior of high- and low-transforming clinical strains during the course of infection, we performed time-lapse microscopy over 12 h of endogenously tagged GFP-LAMP1 HeLa cell line infected by clinical strains. As *S*. Enteritidis is the strain with the strongest epidemiological association with colon cancer (Mughini-Gras et al., 2018), we focused on the 4 *S*. Enteritidis strains previously selected for the transcriptomic analysis (2 low and 2 high transformers) and included the reference strain. All strains were transformed to constitutively express dsRed. We observed that both high-transforming clinical *S*. Enteritidis strains are capable of replicating in a LAMP1-positive SCV, forming SIFs and hyper-replicating in the host cytosol of HeLa cells (Figure 6A, <u>Movie 1</u>). Besides, the replication rate of both high-transforming strains was strikingly higher than the reference strain (Figure 6B). While the two low-transforming *S*. Enteritidis strains could also form SIFs and occasionally hyper-replicate in the cytosol, the number of infected cells and the pace of intracellular replication were significantly lower (Figure 6A-B, <u>Movie 1</u>). We repeated these experiments in MEF Arf^-/-^ cMYC^+^, the cell line used for the soft agar transformation experiments. While all strains had a lower replication rate in MEFs than in HeLa cells, we still observed higher and lower virulence for the high- and low-transforming strains, respectively (Figure 6C, <u>Movie 2</u>). Of note, we observed that both *S*. Typhimurium and *S*. Enteritidis could spontaneously hyper-replicate in MEFs (<u>Movie 2</u>), complementing a previous observation that *S*. Typhimurium deleted for the effector SifA could hyper-replicate in MEF (Thurston et al., 2016).

**Figure 6:**
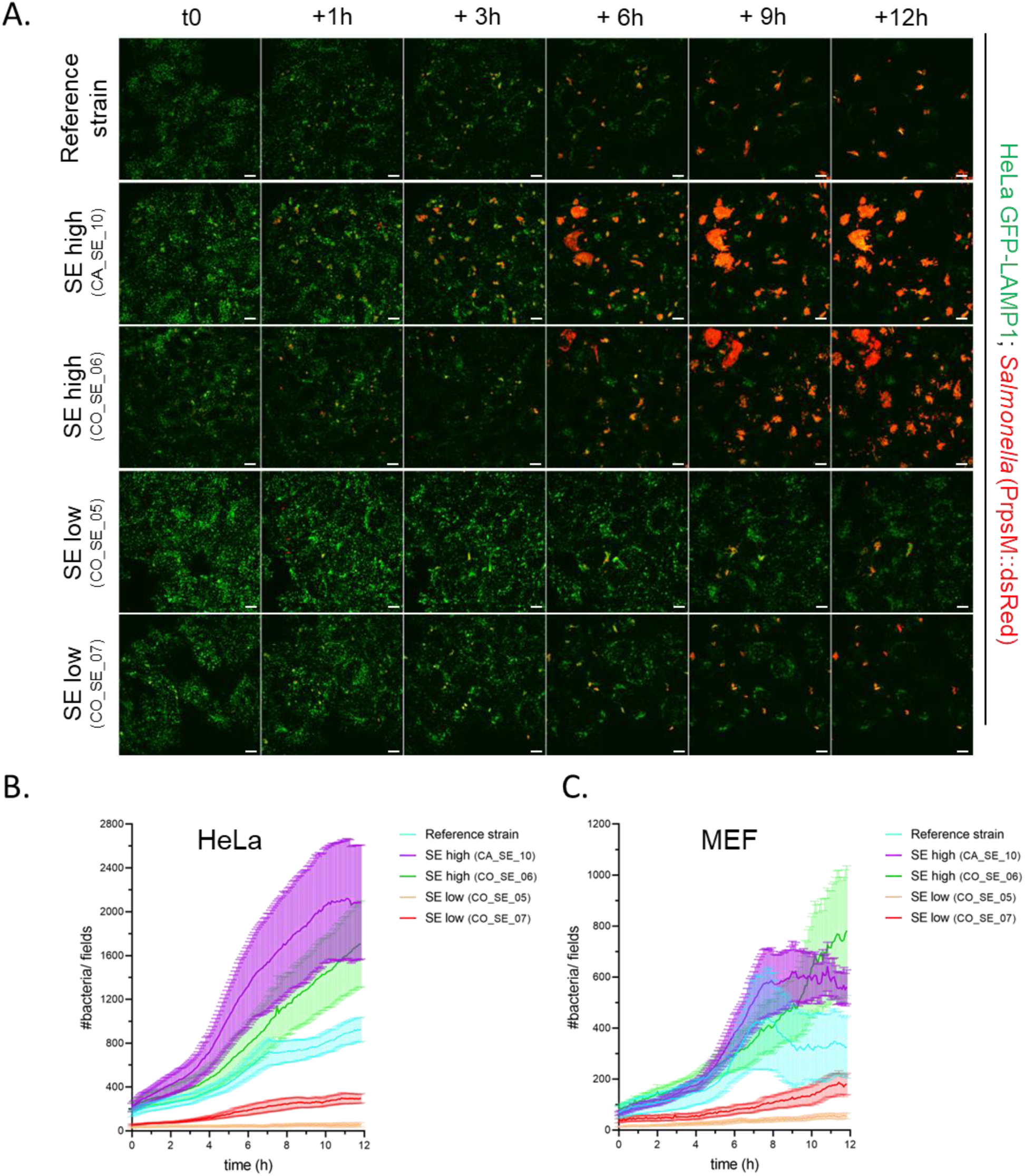
High NTS transformation efficiency correlates with high intracellular growth. A. Acquisition by live microscopy of HeLa cell infections by different *Salmonella* strains: reference strain, CA_SE_10 (SE high), CO_SE_06 (SE high), CO_SE_05 (SE low), and CO_SE_07 (SE low). HeLa cells endogenously express GFP-LAMP1. *Salmonella* strains constitutively express dsRed. Scale-bar: 10 µm. For 5 min time-lapse acquisition, see Movie 1. B-C. Number of bacteria per field along 12 h acquisition of infected HeLa cells (B) or MEF (C). *- back to text.*

To test whether the different strains use a vacuolar high SPI2-expressing lifestyle, or a cytosolic low SPI2-expressing lifestyle (Powers et al., 2021), we designed a SPI2 reporter plasmid where GFP is expressed under the control of the SPI2 promoter *PsseA*, and dsRed is constitutively expressed under the promoter *PrpsM* (named pFPV25-cR-2iG, Figure 7A). We transformed the four clinical strains and the reference strain with this plasmid and followed infections of HeLa cells and MEFs for 12 h by time-lapse confocal microscopy. We observed that all strains start expressing SPI2 within the first hours of the infection in both HeLa (Figure 7B, <u>Movie 3</u>) and MEFs (Figure S6, <u>Movie 4</u>), although hyper-replicating bacteria remained SPI2-negative, similar to what was previously reported for the lab strain *S*. Typhimurium SL1344 (Knodler et al., 2014). When analyzing the rate of bacteria expressing GFP over dsRed during the course of the infection, the high-transforming *S*. Enteritidis strains had a 2.1 and 2.9 times faster SPI2 induction than the reference strain in HeLa cells and MEF, respectively (Figure 7C-D). While the fast induction of SPI2 may explain the higher intracellular fitness of the high-transforming strain, the bacterial capacity to also hyper-replicate in the cytosol in a SPI2 independent manner illustrates the high adaptative growth capacity of these strains in the intracellular environment. Altogether, these results support an increased virulence of high-transforming clinical NTS strains.

**Figure 7:**
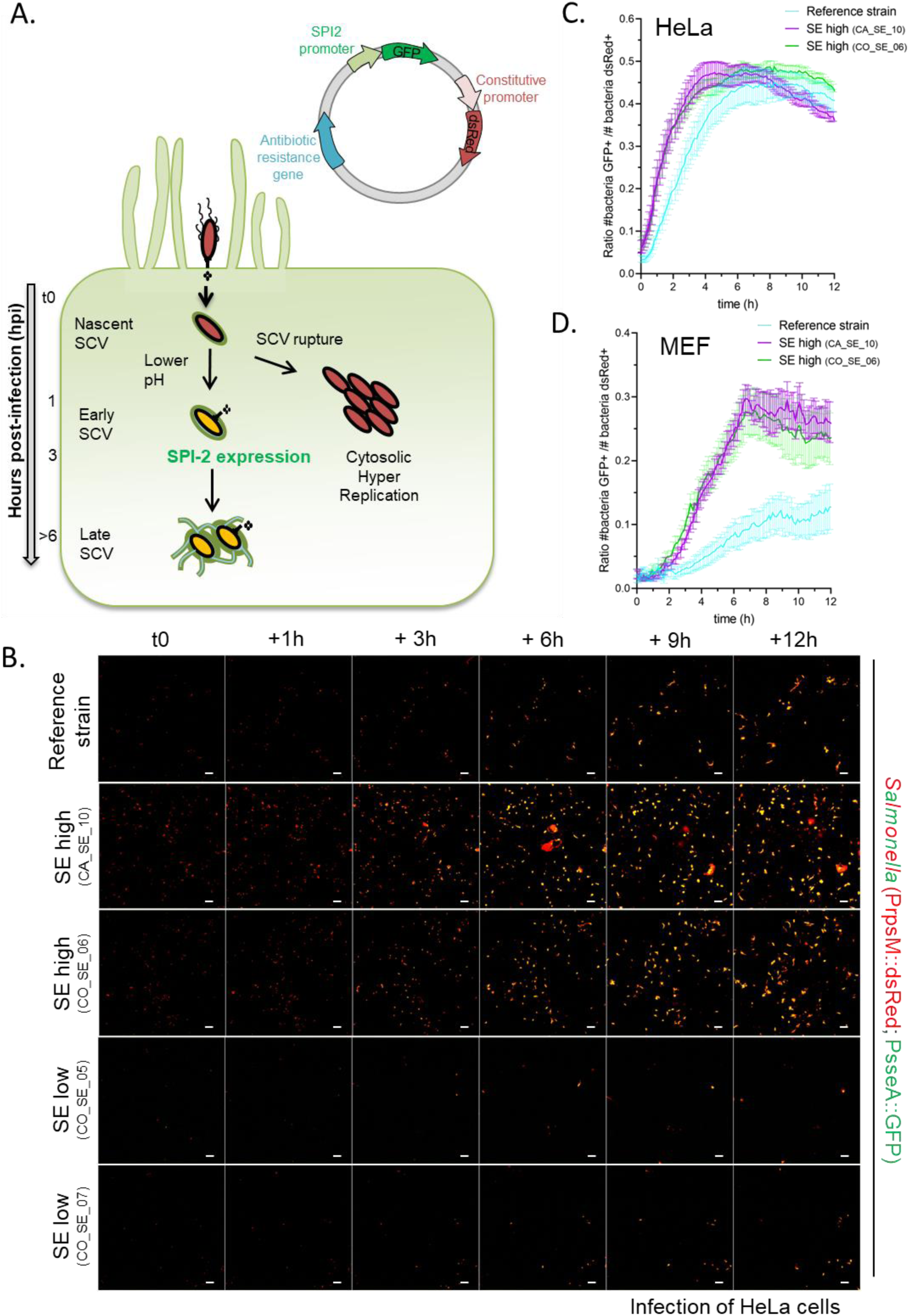
The enhanced intracellular fitness of high-transforming strains is observed in both vacuolar and cytosolic niches. A. Schematic of SPI2 expression reporter system. The red fluorophore dsRed is constitutively expressed using the *PrpsM* promoter while the green fluorophore GFP is expressed upon SPI2-induction using the SPI2 promoter PsseA. SPI2 induction requires SCV acidification and is thus associated with intravacuolar salmonellae. B. Acquisition by live microscopy of HeLa cells infected by different *Salmonella* strains: reference strain, CA_SE_10 (SE high), CO_SE_06 (SE-high), CO_SE_05 (SE low), and CO_SE_07 (SE low). *Salmonella* strains constitutively express dsRed and express GFP upon SPI2 induction. Scale-bar: 20 µm. For DIC acquisition and 5 min time-lapse acquisitions, see Movie 3. C-D. *Salmonella* SPI2 induction during HeLa cell (C) or MEF (D). 3 fields of view per condition, experiment performed in triplicate. *- back to text.*

## DISCUSSION

Upon infection by intracellular pathogens, numerous biochemical mechanisms of the host cell are hijacked and rewired to either promote the pathogen’s survival and replication or limit host defense mechanisms. When the host cells survive such invasions, they are likely to keep a trace of these encounters. As a result, long-term side effects on the host can emerge after the acute infection phase and its recovery. Many “non-infectious” diseases, such as cancer, neurodegenerative disease, autoimmune disease, and very recently endometriosis, are now considered in light of previous infection events (van Elsland and Neefjes, 2018; Backhurst and Funk, 2023; Kivity et al., 2009; Muraoka et al., 2023). Studying these long-term effects is technically challenging due to their low occurrence and long delay after infection. Here, we combined two nationwide patient registries to retrieve stored NTS strains isolated from patients >1 year before colon cancer development. Pathology of colon tumors from patients either with a history of a severe NTS infection or not showed a different grading. This observation suggests that NTS infection may support a different colon cancer program than other tumor-inducing factors.

The clinical outcomes are dependent on host parameters (immune system, pre-existing colon mutations, etc.) that may counteract or synergize with the clinical NTS strain intrinsic transformation efficiency (Lee-six et al., 2019). To extract the potentially “dangerous” NTS strains from the many factors that may contribute to colon cancer formation, we measured the transformation efficiency of the clinical strains using an *in cellulo* assay, eliminating many variable host factors, and used these intrinsic transformation efficiency values for in-depth characterization of NTS features correlating with transformation. While we observed that cancer-associated NTS can transform cells *in cellulo,* we also noticed that a fraction of the control NTS strains isolated had a high transformation efficiency *in cellulo,* suggesting that cell transformation is a widely spread attribute of clinical NTS. Despite the multifactorial etiology of cancer with the cumulative influence of environmental and genetic carcinogenic factors, we could observe a correlation between *in cellulo* assessment of NTS strain transformation efficiency and clinical association as the case strains had higher transformation efficiency than their matching control strains.

Our study reveals that the strain attachment and invasion efficiency in polarized Caco2 cells, and invasion efficiency in MEF are positively correlated with the transformation efficiency, although they could not completely explain the variation herein. Therefore, the success of bacteria in transforming eukaryotic cells is partly dependent on their ability to attach and invade these cells. As *Salmonella* has to infect a pre-transformed cell in order to induce transformation (Scanu et al., 2015), an increased invasion capacity is likely to foster the chance for the bacteria to encounter pre-transformed cells in the gut and promote cancer.

We used the transformation efficiency measured for each strain to test for correlating genomic and transcriptomic features. While there were very few and inconclusive genetic features whose presence/absence was associated with transformation efficiency, the transcriptomic analysis identified several hundred genetic features with differential expression across the spectrum of transformation efficiency. These results suggest that the transformation efficiency of NTS is not necessarily defined by the genetic makeup of the bacteria but by the differential expression of particular genes. Previous research has also shown that even though related bacterial strains have similar gene contents, transcription factor binding sites can be easily gained or lost, with subsequent impact on the transcription patterns (Perez and Groisman, 2009). Although we could not identify a clear-cut separation of pathways enriched in the two categories of transformation efficiency, there appeared to be entire operons differentially expressed between them. Thus, in the SPI2-inducing condition, high-transforming strains seem to overexpress genes associated with flagellar assembly, chemotaxis, energy metabolism, and signal transduction, while the low-transforming strains, under the same experimental conditions, appear to overexpress genes related to quorum sensing. Flagellar assembly has been suggested to be coupled with the progression through the cellular cycle and the formation of a differentiated daughter cell (Mangan et al., 1999). This correlates with our finding that high NTS transformation efficiency is accompanied by high intracellular growth of the bacteria. Additionally, a high level of oxidative phosphorylation and ATP synthesis is consistent with the higher need for energy of an increasing bacterial population. One of the few pathways that appeared to be enriched only in low transformers – that of quorum sensing – contains genes that are also linked to signal transduction and that promote intramacrophage multiplication of *S*. Typhimurium (Gannoun-Zaki et al., 2014); this set of signal transduction genes is smaller and different from the one in high transformers and possibly with lower efficiency in bacterial growth. This suggests that the bacterial strains with differential transformation efficiencies use preferentially alternative signaling pathways. Aligned to this, we found that in the SPI2-inducing condition, the high-transforming strains expressed more genes belonging to specific regulons than the low-transforming strains. This result indicates that the high transformers may be more sensitive to environmental cues present in the SPI2-inducing condition that are detected by regulatory proteins. This hypothesis is fostered by our microscopy observations of faster induction of SPI2 in the high-transforming strains.

Whether the differential expression of some bacterial genes is directly involved in promoting mammalian cell transformation is unclear. A technical difficulty behind this objective is that we did not identify a specific pathway that was activated, but rather many pathways associated with the observed phenotype of high metabolic flexibility and intracellular fitness. Based on our results, we can propose that *Salmonella*’s metabolic flexibility and intracellular fitness may induce host cell transformation, possibly due to the host metabolic shift generated by *Salmonella* growth.

Non-mutually exclusive models suggest the involvement of the *Salmonella* effectors SopB and AvrA, coinciding with activated host Akt and JNK pathways (Scanu et al., 2015; Lu et al., 2016). Of note, both Akt and JNK pathways are also involved in metabolic reprogramming and cancer metabolism (Papa et al., 2019; Hoxhaj and Manning, 2020), suggesting a cumulative effect of *Salmonella* hijacking mechanisms pushing for metabolic reprogramming and cell transformation. Besides, we previously showed that the overexpression of the proto-oncogene cMYC was required to observe fibroblast transformation after *Salmonella* infection, using an anchorage-independent growth assay (Scanu et al., 2015), and cMYC is also one of the key players regulating cancer cell metabolism (Dong et al., 2020). Finally, *Salmonella* tropism towards (pre-)transformed cells suggests a cumulative effect during repetitive exposure (Forbes et al., 2003; van Elsland et al., 2022; Aganja et al., 2022). Thus, repetitive pressure on cell metabolic equilibrium could push the cell toward transformation. This model suits the observation that we did not identify genetic alterations in *Salmonella*-transformed MEF (Scanu et al., 2015).

Altogether, this multi-faceted explorative study based on information obtained in a nationwide epidemiological study identified novel features of NTS linked to colon cancer and cell transformation. In particular, the increased attachment, invasion, and intracellular fitness observed in the high-transforming strains reveal an association between virulence and transformation efficiency and open avenues for future novel investigations linking infection, metabolic changes, and cell transformation. Thus, circulating NTS strains display different phenotypes linked with different potential for host cell transformation.

## ACKNOWLEDGMENTS

This work was supported by a KWF grant ‘‘Bacterial food poisoning and colon cancer; a cell biological and epidemiological study’’ (2017-1-11001) (to J.N., E.F., and L.M.-G.), an ERC Advanced grant ERCOPE (to J.N.), and the Netherlands Organisation for Health Research and Development (ZonMw) (to J.N., E.F., and L.M.-G.) (grant number 522004001). V.S. was supported by the LUMC MSCA-IF Seal of Excellence program.

## AUTHOR CONTRIBUTIONS

Histology staining and analysis: A.N-B; Transformation and invasion efficiency: V.S., D.v.E, L.V.; Gastrointestinal tract model system: A.v.H; Genomic analyses and Genome-wide association analysis: C.C, J.D., Transcriptomic analyses: V.S., C.C.; Metabolic profiling: C.C, J.D, A.v.H; Intracellular growth analyses: V.S.; Writing – Original Draft, V.S., C.C, J.D., D.v.E; Writing – Review & Editing, V.S., C.C., L.M-G. and J.N.; Tools development: L.J. and J.A.; Funding Acquisition, L.M.-G., E.E., E.F., and J.N.

## METHODS

### EXPERIMENTAL MODEL AND SUBJECT DETAILS

#### Bacterial strains and plasmids

*S*. Typhimurium strain SL1344 was used as the reference strain in the *in cellulo* invasion and transformation assays. The 60 clinical NTS strains (listed in Supplementary Table S1) were obtained from the bacterial strain collection of the RIVM as described above. These strains were identified during routine diagnostic activities on patients with gastroenteritis between 1999 and 2015 by medical microbiology laboratories in the Netherlands, and subsequently sent to the RIVM for further typing as part of the national laboratory surveillance system for *Salmonella*. All strains were confirmed to be gentamicin-sensitive.

To follow bacterial growth, the bacteria were transformed with the pGG2 plasmid (Lelouard et al., 2010) for constitutive expression of dsRed under the promoter *PrpsM*. To follow SPI2 expression, a variant of the pGG2 plasmid was created by inserting GFP preceded by the SPI2 promoter Pssea. Of note, it was necessary to use a 1.5 kb region upstream of sseA as promoter to observe SPI2 induction in NTS. This region was amplified by PCR using the oligonucleotides PsseA fwd Asp718 (5′ cccaggtacccgtattcttgattttcatcggtgg 3′) and PsseA rev XbaI (5′ cccatctagatgccctttcagcaagctgttgac 3′), as previously described by Brown et al., 2005. Besides the ampicillin resistance cassette was replaced by kanamycin. We named this plasmid pFPV25-cR-2iG (for backbone name, <u>c</u>onstitutive ds<u>R</u>ed expression and SPI<u>2</u>-induced <u>G</u>FP expression, Addgenes #204618).

#### Mammalian cell lines

MEFs, WT HeLa, and endogenously GFP-tagged LAMP1 HeLa cells were cultured in Dulbecco’s modified Eagle’s medium (DMEM) supplemented with 8% Fetal Bovine Serum (FBS) at 37°C, 5% CO2. MEFs *Arf*-deficient and overexpressing c-MYC were generated in a previous study (Scanu et al., 2015). LAMP1-mGFP HeLa cells were endogenously tagged using CRISPR-CAS9 technology (this study). Briefly, a homologous repair construct was made by cloning 300 bp genomic regions up and downstream of the stop residing on the last exon from LAMP1 into mGFP-T2A-Puromycin construct. gRNAs were designed using the CRISPOR online tool (Concordet and Haeussler, 2018) and cloned into pX330 U6 hSpCas9 vector using the following cloning primers: NdeI cEndo LAMP1 5 fw (actacgcatatgtctctgcagcggcttccc); NheI cEndo LAMP1 5 rv (actacggctagcgatagtctggtagcctgcgtgac); BglII cEndo LAMP1 3 fw (actacgagatcttagcctggtgcacgcagg); SalI cEndo LAMP1 3 rv (actacggtcgacaccccctcagagagagcacc); cEndo LAMP1 gRNA S:95 fw (caccgctatctagcctggtgcacgc) and cEndo LAMP1 gRNA S:95 rv (aaacgcgtgcaccaggctagatagc). Homologous repair and pX330 constructs were cotransfected into Hela cells using XtremeGene. The transfected cells were expanded over 8 days. Next, GFP-positive cells were single-cell sorted with flow cytometry. The newly generated single-cell clones were validated by Western Blot and microscopy. All cell lines used in the study were regularly checked for mycoplasma.

### METHOD DETAILS

#### Histology staining and analysis

Formalin-fixed paraffin-embedded (FFPE) tissue blocks of 24 patients with proximal CC with a registered NTS infection in the past and 67 tissue blocks from age- and gender-matched controls (with proximal CC) without such history were obtained from the Dutch nationwide network and registry of histo- and cytopathology (PALGA). Tissue blocks were requested from different Dutch pathology labs and collected at AmsterdamUMC location VUMC. The tissue blocks were sectioned and the sections were H&E stained according to standard protocols. In addition, the tumor grading was determined for each of the 91 CC patients by an experienced CC pathologist (A. N-B.) who assessed the samples in a blinded fashion. The sample identity was with L.M. at the Dutch Health Institutes and was opened after the completion of the pathology analyses.

#### Invasion and transformation assays in MEF

The step-by-step protocol describing NTS infection of MEFs and anchorage-independent growth assay was recently published (Stévenin and Neefjes, 2023). Briefly, NTS were grown overnight at 37°C in LB medium one day prior to the infection. On the day of the infection, the bacterial overnight cultures were diluted at 1:33 in LB and the subcultures were incubated for 2-3 h at 37°C in a rotating incubator (220 rpm). Cells were infected with NTS at a multiplicity of infection (MOI) 20 in DMEM medium without antibiotics for 20 min at 37°C, 5% CO_2_. The samples were washed 3 times to eliminate extracellular bacteria and incubated in DMEM supplemented with 8% FBS and 100 μg/mL gentamicin for 1 h to kill the remaining attached extracellular bacteria.

For CFU assays, cells were washed and lysed with lysis buffer (ddH2O + 1%NP-40). Serial dilutions of the lysate were plated on LB plates and incubated at 37°C overnight. CFUs were counted the next day.

For soft agar anchorage-independent growth assays, a bottom layer of soft agar consisting of 0.7% low melting point agarose in DMEM supplemented with 8% FBS and 10 μg/mL gentamicin was prepared during the subculture time (Stévenin and Neefjes, 2023). After 1 h incubation with gentamicin at 100 μg/mL, the MEF medium was changed for DMEM supplemented with 8% FBS, and 10 μg/mL gentamicin and MEF were grown for another 2 h at 37°C, 5% CO_2_. Then, MEFs were trypsinized, collected, and resuspended at 25,000 cells/mL in DMEM medium supplemented with 8% FBS, 10 μg/mL gentamicin, and 0.35% low melting point agarose (UltraPure™, Invitrogen). This cell-containing solution was poured on the soft agar bottom layer to form the soft agar top layer. Soft-agar cultures were incubated for 2-3 weeks at 37°C, 5% CO2. The number of cell colonies was assessed by using GelCountTM (Oxford Optronix, UK).

#### Gastrointestinal tract model system

The NTS strains were cultured overnight at 37°C and subsequently exposed to conditions resembling the human digestive tract in a gastrointestinal tract (GIT) model system (Oliveira et al., 2011; Wijnands et al., 2017; Kuijpers et al., 2019). An overnight culture (ON) of each clinical NTS strain was sequentially exposed to simulated gastric fluid (SGF) and simulated intestinal fluid (SIF) at 37°C for 30 min and 2 h, respectively. After that, differentiated Caco-2 cells mimicking the small intestinal epithelium were inoculated with the SGF/SIF/bacterial mixture at 37 °C for 1 hour on a 12-well plate to test the bacterial attachment (ATT) and invasion (INV) capacities. Between each step of the GIT model (ON, SGF, SIF, ATT, and INV), serial 10-fold dilutions of samples were made and NTS bacteria present were enumerated. For quantification of attachment, 6 out of 12 wells containing the Caco-2 cells were lysed (to release attached and invaded bacteria), whereas, for enumeration of invaded bacteria only, the other 6 out of the 12 wells were treated with gentamicin to kill attached bacteria before lysing cells to release invaded NTS. Details about the cell cultures, the dilution steps, the compositions of SGF and SIF, and the experimental procedures are described elsewhere (Oliveira et al., 2011; Wijnands et al., 2017; Kuijpers et al., 2019).

#### Genomic analyses

All 60 clinical strains were submitted to whole-genome sequencing (WGS). DNA isolation, 2×125 bp paired-end library preparation and WGS analysis on a HiSeq 2500 platform (Illumina) was performed by BaseClear (Leiden, the Netherlands). All resulting fastq files were subjected to quality control with CheckM v1.0.7 (Parks et al., 2015), and *de novo* assembled using an in-house developed pipeline (https://github.com/RIVM-bioinformatics/Juno_pipeline). The assembled genomes were analyzed with the SISTR application to confirm the *Salmonella* serovar (Yoshida et al., 2016). The assembled genomes were screened for the presence of 233 *Salmonella* virulence factors (Kuijpers et al., 2019). Genome annotation of the assembled genomes was performed with Prokka v1.14.6 (Seemann., 2014). Next, the annotation data was used as input for Roary v3.13.0 to construct the core- and accessory genome of the 60 clinical strains, with a blastp identity cut-off of 95%, indicating a presence in at least 57 out of the 60 clinical NTS strains for a gene to be defined as part of the core-genome (Page et al., 2015). The core genome alignment from Roary was used to build a phylogenetic tree in RAxML v8.2.12 (Stamatakis., 2014). Protein functional annotation was performed with Pannzer2 (Törönen et al., 2018).

#### Genome-wide association analysis (GWAS)

The genes in the pangenome of NTS as inferred by Prokka and Roary were used to inform the association tests performed by TreeWAS. Several genome-wide association tests were conducted using the R-package TreeWAS, which accounts for recombination and population structure (Collins et al., 2018). Reconstruction of ancestral states was done with the parsimony method, and phylogenetic trees were constructed using maximum likelihood (ML) phylogeny. For all association tests, a false discovery rate (FDR) correction was applied to account for multiple testing. The outcome variable used in the association tests was the ranked transformation capacity of the bacterial strains, as inferred from the *in vitro* tests (see above). Statistical analyses were performed with R v3.6.2.

#### Transcriptomic analyses

For *in vitro* stimulation of SPI1 operons, bacteria were grown overnight in 5 mL Lennox Broth, diluted 1:1000 into 100 mL Lennox Broth, grown at 37°C in a 500 mL Erlenmeyer flask in a rotating incubator (220 rpm) until early stationary phase, corresponding to an OD_600_ of 2.0 (Kröger et al, 2013). For *in vitro* stimulation of SPI2 operons, bacteria were grown overnight in 5 mL Lennox Broth, and diluted 1:50 into 100 mL of acidic low-phosphate growth medium (PCN medium at pH 5.8, containing 80 mM MES, 4 mM Tricine, 100 µM FeCl_3_, 376 µM K_2_SO_4_, 50 mM NaCl, 0.4 mM K_2_HPO_4_/KH_2_PO_4_, 0.4 % glucose, mM NH_4_Cl, 1 mM MgSO_4_, 0.01 mM CaCl_2_, 10 nM Na_2_MoO_4_, 10 nM Na_2_SeO_3_, 4 nM H_3_BO_3_, 300 nM CoCl_2_, 100 nM CuSO_4_, 800 nM MnCl_2_, 1 nM ZnSO_4_) until reaching an OD_600_ of 0.3 (Lober et al, 2006). At the desired OD, the bacterial cultures were centrifuged at 4200 *g* for 10 min at 4°C. Pellets were resuspended in ShieldBuffer (BaseClear) and transferred to a transport tube.

The RNA extraction, rRNA depletion using the RZP (Illumina) kit, library preparation, and sequencing were performed by the external company BaseClear. 250 ng total RNA in RZP was used for sequencing. Single-end or paired-end sequence reads were generated using the Illumina NovaSeq 6000 or MiSeq system. The sequences generated with the MiSeq and NovaSeq 6000 sequencers were performed under accreditation according to the scope of BaseClear B.V. (L457; NEN-EN-ISO/IEC 17025). Fastq sequence files were generated using bcl2fastq version 2.20 (Illumina). The samples went through two steps of quality assessment. The initial quality assessment was based on data passing the Illumina Chastity filtering. Subsequently, reads containing the PhiX control signal were removed using an in-house filtering protocol. In addition, reads containing (partial) adapters were clipped (up to a minimum read length of 50 bp). The second quality assessment was based on the remaining reads using the FastQC quality control tool version 0.11.8.

Based on the pangenome (see section Genomic analyses) we have selected from NCBI a reference sequence with minimum distance to our bacterial strains of interest – *Salmonella* Typhimurium SL1344. The reference genome was annotated for the presence of pathogenicity islands (coined as “regulons” in Avital et al., 2017, and Westermann et al., 2016) using blast 2.10.1 against the PAIDB database (Yoon et al., 2015; accessed 05 July 2021), and for putative operons using the Operon-mapper web server (Taboada et al., 2018). Mapping of the reads to the reference genome was performed with Kallisto v 0.46.2 (Bray et al., 2016) and subsequent filtering of the reads mapping to various species of RNA (ribosomal, transfer, and transfer-messenger) was performed in R v>4.1. The various bacterial strains showed differences in their genomes (see Genomic analyses) with some genomic loci being present in only a fraction of the strains. To reduce a possible bias in differential gene expression, we have filtered the genomic loci to include only the ones present in all bacterial strains. The differential gene expressions were analyzed in the R package *DSeq2* v 1.40 using three subsets of the RNAseq sequences: SPI2 vs SPI1, low transformers vs high transformers for each experimental condition (SPI1- and SPI2-induction). To compare our results with earlier research on intracellular behaviors of *Salmonella spp*. We have included in our analyses not only the genome annotation based on Prokka and pathways from KEGG but also the custom annotation of regulatory pathways described in Powers et al., 2021.

#### Metabolic profiling

The utilization of carbon (C), nitrogen (N), phosphorus (P), and sulfur (S) sources by the 60 clinical NTS strains were analyzed using the BioLog^®^ Phenotype MicroArray (plates PM1, PM3, and PM4), which allows for high-throughput metabolic quantification of bacterial respiration and growth on a range of different substrates (Shea et al., 2012). 95 carbon (PM1), 95 nitrogen (PM3), 59 phosphorus (PM4) and 35 sulfur (PM4) sources were tested (Supplementary Table S7). Briefly, the quantification is based on redox technology, in which cell respiration is measured by the degree of irreversible reduction of a tetrazolium dye. Following Biolog instructions, a cell suspension of each individual cultured NTS strain and a defined medium (including a dye) were added to 96-well plates containing a single C-, N-, P-, or S-source in each well. Plates were incubated at 37°C for 24h and color formation was measured every 15 min using an ELx808 Microplate Reader and Gen5 software (BioTek).

#### Intracellular growth analyses

Cells were grown in a 4-chamber Ibidi glass-bottom dish. The day before the infection, bacteria were grown overnight in 5 mL Lysogeny Broth (LB) medium at 37°C in an orbital shaker (220 rpm). Growth media was supplemented with 100 mg/mL ampicillin or 25 mg/mL kanamycin depending on the resistance cassette of the plasmids (pGG2 and pFPV25-cR-2iG, respectively). On the day of the infection, bacteria were subcultured at a 1:33 dilution in LB and incubated at 37°C for 3 h (i.e., until the late exponential phase). Bacteria were then washed once and resuspended in EM buffer (120 mM NaCl, 7 mM KCl, 1.8 mM CaCl_2_, 0.8 mM Mg Cl_2_, 5 mM glucose, 25 mM HEPES, pH 7.3). Bacterial concentration was determined using OD600, and bacteria were diluted to the desired multiplicity of infection (MOI) in warm EM buffer. Bacteria were added to the cells and incubated for 30 min at 37°C, 5% CO_2_. Cells were gently washed 2-3 times with EM buffer to eliminate extracellular bacteria. The cell medium was changed for EM buffer supplemented with 10% FBS and gentamicin at 20 mg/mL and the cells were placed in a microscope chamber at 37°C, 5% CO_2_. Confocal microscopy was performed on an Andor Dragonfly 200 spinning disk microscope equipped with 40x/1.30 oil HCX Plan Apo objective, 63x/1.40-0.60 oil Plan Apo objective, and an Andor Zyla 4.2 PLUS sCMOS Camera. 3 fields of view were selected per conditions and all conditions were acquired in parallel every 5 min for 12 h. Z-stacks of 5.5 µm with 0.3 µm steps were acquired in both the GFP (laser 488 nm) and dsRed (laser 561 nm) channels. Maximum z-projections were performed on all movies and the number of positive dsRed pixels (above noise threshold) was measured over time using ImageJ v1.5. The number of bacteria was estimated by dividing the number of positive pixels by the average pixel size of one bacterial cell (estimated over 100 bacteria). To quantify SPI2 induction, a binary image was created on ImageJ using the dsRed channel to detect the position of the bacteria at each time frame and the number of GFP-positive bacteria per field was divided by the number of dsRed-positive bacteria. The strains “SE low CO_SE_05” and “SE low CO_SE_07” were excluded from this analysis as the too-low number of bacteria per field generated a high variability and negative bias in the #bacteria GFP+ /#bacteria dsRed+ ratio.

### QUANTIFICATION AND STATISTICAL ANALYSIS

#### Statistical analysis of NTS infection and transformation efficiency

Each strain infection efficiency and transformation efficiency were measured independently 2 times (biological replicate). All experiments included technical triplicate and an internal control using the reference strain SL1344 for normalization. In the few cases where strong variation was observed between biological duplicates (n=4), a third repetition was performed and the average of the 3 experiments was used. The relation between the transformation efficiency and the invasion efficiency measured either on cell cultures or in the GIT system was quantified using orthogonal regressions and Pearson correlation coefficients.

#### Transcriptomic analyses

15 different experimental conditions (8 strains grew within 2 different growth media) were compared. For each condition, sample preparation, RNA extraction, and sequencing were performed at least 3 times. However, the quality of the RNA sequences was heterogeneous between experimental conditions. Upon the SPI1-induced condition, we obtained 16 biological replicates of seven bacterial strains, while upon induction of the SPI2-induced condition, we had 28 biological replicates of eight bacterial strains.

#### Phenotype microarray

The analyses were performed twice for each of the 60 NTS strains. The ratio of the integrals of each C-, N-, P-, and S-source and a negative control (i.e. the PM1, PM3, and PM4 plates contain a negative control well without substrate for each source type), was used as the outcome for further analysis. Hierarchical cluster analysis (HCA) and principal component analysis (PCA) were performed on the scaled data for each of the three plates to compare the metabolic phenotypes of NTS strains obtained from cases versus controls taking into account the transformation efficiency of the strains. We defined the optimal clustering method for hierarchical clustering and the average linkage clustering fit best to the data (lowest Gower distance and highest cophenetic correlation). Spearman’s rank correlation test followed by Bonferroni correction were used for multiple testing of correlations between substrate utilization and transformation efficiency. The analyses were performed in R version 1.4 1103.

#### Intracellular growth analyses

The number of bacteria per field is estimated at every time point by measuring the number of dsRed positive pixels per field (above background level) divided by the average pixel size of one bacterium (measured over 100 bacteria). To calculate the ratio of GFP-positive over dsRed-positive bacteria, first, all bacteria were detected using the dsRed channel in Fiji. Then the dsRed-positive bacteria were used as a mask to detect the GFP-positive bacteria. This allowed to detect only the bacterial GFP signal and avoid detecting the GFP signal from the MEF nuclei. The ratio of GFP-positive over dsRed-positive bacteria was calculated per field at each time point. Graphs in Figures 6 and 7 display the mean and SEM of 9 fields of view per condition, measured over 3 independent experiments.

## FIGURES

**Figure S1:**
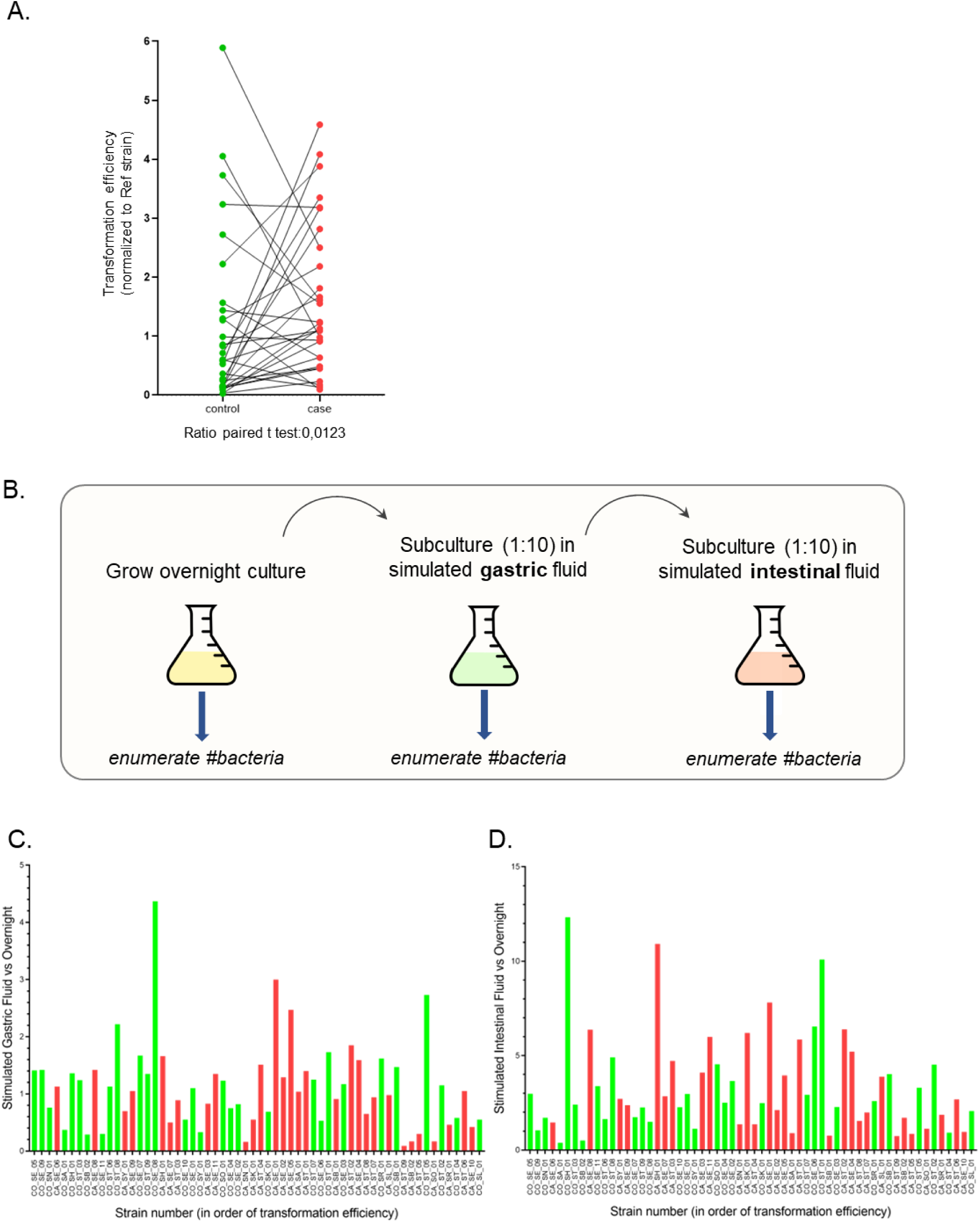
A. Transformation efficiency (mean) of the matching pairs of case-control strains, normalized to the reference strain. B. Schematic representation of the simulated gastrointestinal passage. Overnight bacterial cultures are sequentially enumerated and diluted in simulated gastric fluid and simulated intestinal fluid. C. Ratio of CFU counts after growth in simulated gastric fluid versus overnight culture for each clinical NTS. Strains are ordered by transformation efficiency in the x-axis. D. Ratio of CFU counts after growth in simulated intestinal fluid versus overnight culture for each clinical NTS. Strains are ordered by transformation efficiency in the x-axis. *- back to text.*

**Figure S2:**
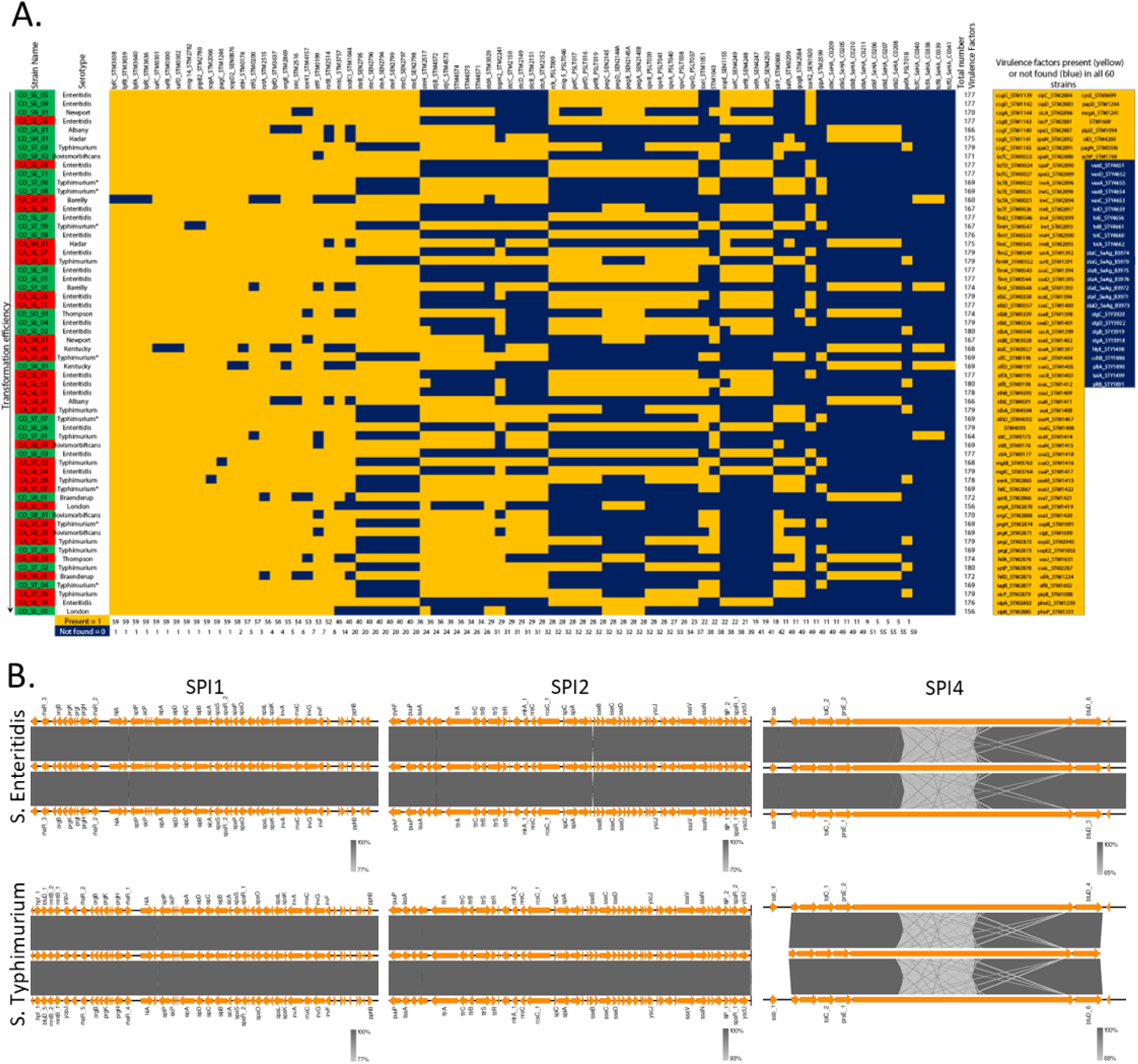
A. Virulence factors present (yellow) or not found (blue) in the 60 NTS strains. *monophasic variant as shown by WGS analysis. [High-resolution panel](https://drive.google.com/file/d/1cdh7Cd66Z_hB03plOx3RuhbYAW135t6E/view?usp=sharing). B. Comparison of *Salmonella* pathogenicity island (SPI) organization between strains. Alignment of SPI1 (encoding T3SS-1), SPI2 (encoding T3SS-2), and SPI4 (encoding T1SS) for *S*. Enteritidis strains CO_SE_05 (transformation efficiency: low; 0.03), CO_SE_06 (transformation efficiency: medium; 1.30), and CA_SE_10 (transformation efficiency: high; 4.59), and S. Typhimurium strains CO_ST_06 (transformation efficiency: low; 0.15), CA_ST_08 (transformation efficiency: medium; 1.81), and CO_ST_04 (transformation efficiency: high; 4.05). Orange arrows: ORF pointing in the direction of the reading frame. The grey scale represents the percentage of similarity between the two adjacent sequences. *- back to text.*

**Figure S3:**
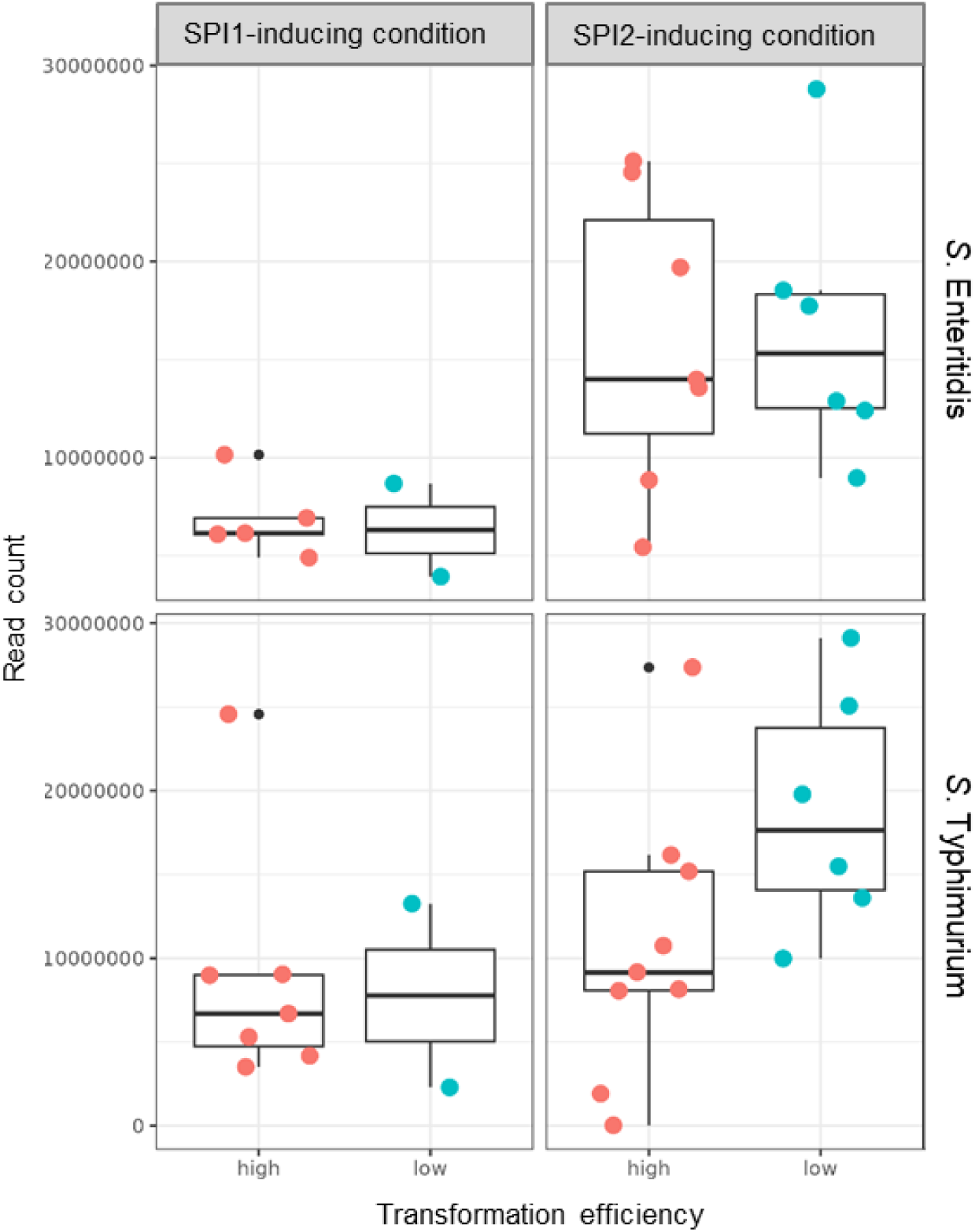
Number of reads identified using the program kallisto matching the coding DNA sequences (CDSs) of the references. *- back to text.*

**Figure S4:**
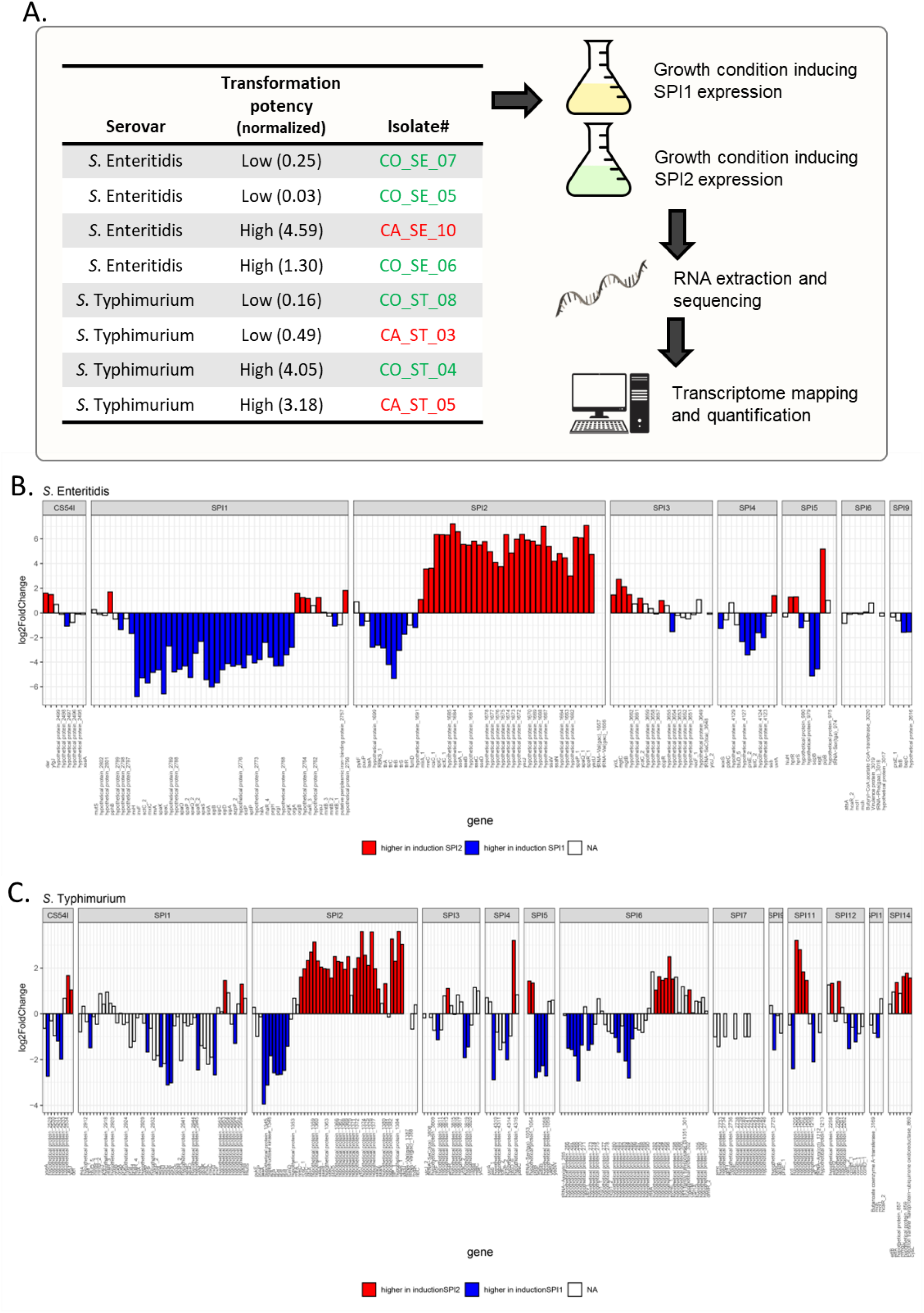
A. Schematic of the transcriptomic analyses pipeline. Eight NTS strains were selected and grown in SPI1- or SPI2-inducing growth medium before harvesting, RNA extraction, sequencing, and transcriptome mapping and quantification. B. Differential gene expression of the genes included in the *Salmonella* pathogenicity islands (SPI) of *S*. Enteritidis clinical strains. *S*. Enteritidis reference CP007507, reads filtered for r−, t−, and tmRNA. The genes with increased expression in the SPI1- and SPI2-inducing condition are depicted below and above the null, respectively. Only the genes that were significantly overexpressed (FDR adjusted p-value < 0.05) are colored. C. Differential gene expression of the genes included in the SPI of *S*. Typhimurium clinical strains. *S*. Typhimurium reference SL1344, reads filtered for r−, t−, and tmRNA. The genes with increased expression in the SPI1- and SPI2-inducing condition are depicted below and above the null, respectively. Only the genes that were significantly overexpressed (FDR adjusted p-value < 0.05) are colored. [high-resolution B-C panels](https://drive.google.com/file/d/1IQSMw4nnff_E_J5gCGzdUmXosInOubiq/view?usp=sharing). *- back to text.*

**Figure S5.**
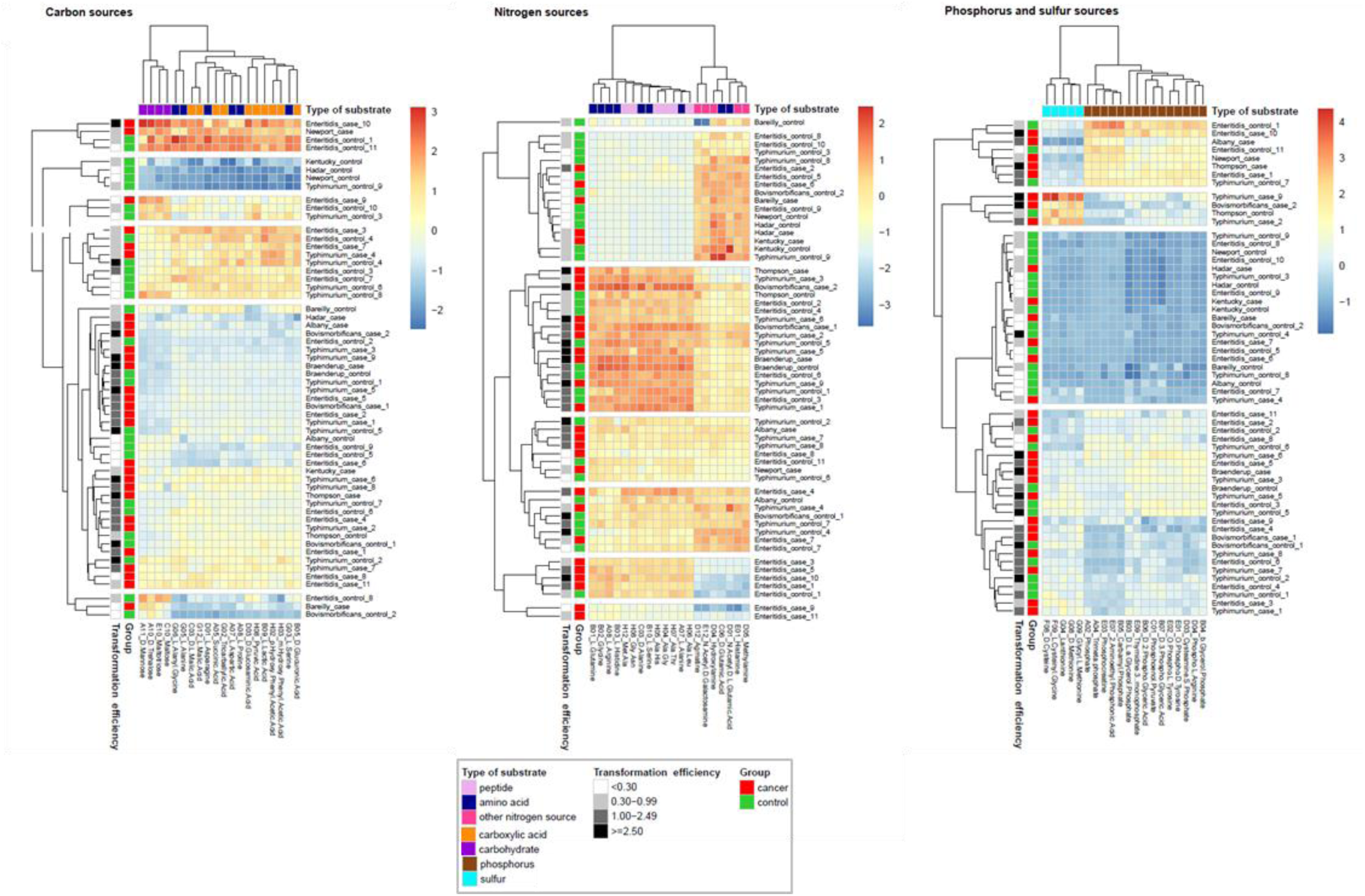
Heatmap of scaled utilization scores of the 60 NTS strains for the top 20 sources mostly contributing to the variance in the carbon, nitrogen, and phosphorus/sulfur data. NTS isolates are clustered based on their utilization scores using the average linkage method. Nutrients are clustered using the average linkage method. *- back to text.*

**Figure S6:**
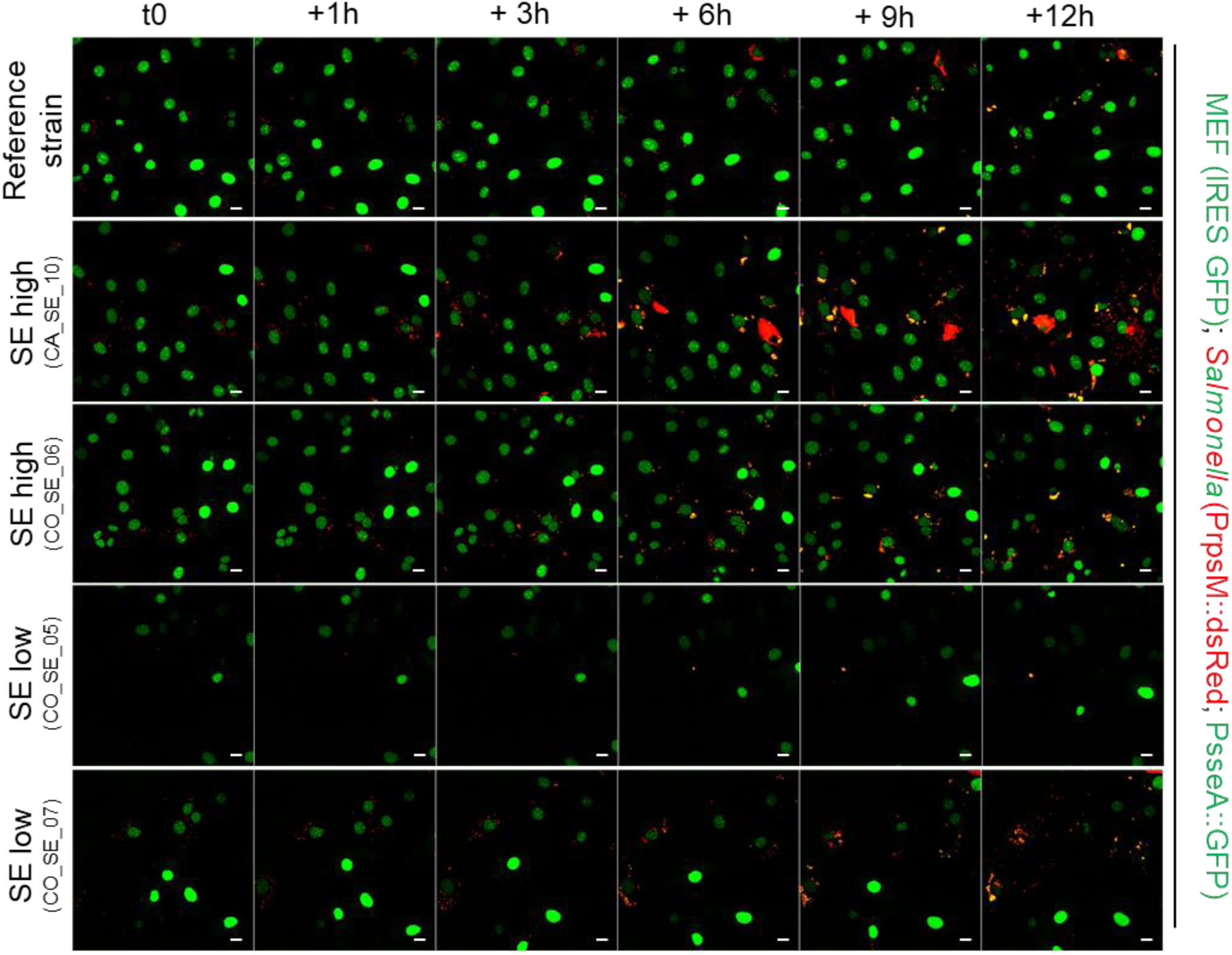
Acquisition by live microscopy of MEF infected by different *Salmonella* strains: reference strain, CA_SE_10 (SE-high), CO_SE_06 (SE-high), CO_SE_05 (SE-low), and CO_SE_07 (SE low). *Salmonella* strains constitutively express dsRed and express GFP upon SPI2 induction. Scale-bar: 20 µm. For 5 min time-lapse acquisitions, see Movie 4. *- back to text.*

## MOVIES

**Movie 1.** Infection of HeLa cells endogenously tagged (GFP-LAMP1) by salmonellae constitutively expressing dsRed. Left panel: Reference strain, middle panel SE high CA_SE_10, right panel: SE low CO_SE_07. Scale bar: 10 µm.

➔ [LINK Movie 1](https://drive.google.com/file/d/1S57-lu0QLRiJULYsdIz3ZXL6JQblPyX7/view?usp=sharing)

➔ <u>back to text</u>.

**Movie 2.** Infection of MEF (Arf-/-; c-MYC+) containing IRES-2NLS-GFP (used for clone selection) by salmonellae constitutively expressing dsRed. Left panel: Reference strain, middle panel SE high CA_SE_10, right panel: SE low CO_SE_07. Scale bar: 10 µm.

➔ [LINK Movie 2](https://drive.google.com/file/d/14GuW71vskPIPPQhluUWGsI-rjcGFckWH/view?usp=sharing)

➔ <u>back to text</u>.

**Movie 3.** Infection of HeLa cells by salmonellae constitutively expressing dsRed and expressing GFP upon SPI2 induction (*PrpsM*::dsRed; *PsseA*::GFP). Left panel: Reference strain, middle panel SE high CA_SE_10, right panel: SE low CO_SE_07. Scale bar: 10 µm. First frame: DIC acquisition before the start of the time-lapse acquisition.

➔ [LINK Movie 3](https://drive.google.com/file/d/153kkDRLfKdGDN7rn57Z3DwXpYwXd9YvV/view?usp=sharing)

➔ <u>back to text</u>.

**Movie 4.** Infection of MEF (Arf-/-; c-MYC+) containing IRES-2NLS-GFP (used for clone selection) by salmonellae constitutively expressing dsRed and expressing GFP upon SPI2 induction (*PrpsM*::dsRed; *PsseA*::GFP). Left panel: Reference strain, middle panel SE high CA_SE_10, right panel: SE low CO_SE_07. Scale bar: 10 µm.

➔ [LINK Movie 4](https://drive.google.com/file/d/1-Zlj52G4OVXGw1e-neisVGaWUQrA6E2y/view?usp=sharing)

➔ <u>back to text</u>.

## TABLES

**Supplementary Table S1.** Characteristics of the NTS infection in individuals who developed colon cancer later in life (i.e. cases) versus those who did not develop cancer (i.e. controls). *monophasic variant as shown by WGS analysis.

➔ [LINK Table S1](https://docs.google.com/spreadsheets/d/1Qp3PIQuA_pDUKxSJCTGldCvOWnvQlrBt/edit?usp=drive_link&ouid=113479068300354772155&rtpof=true&sd=true)

➔ <u>back to text</u>.

**Supplementary Table S2.** Strain invasion and transformation efficiency in MEFs, normalized by the invasion efficiency of the reference strain.

➔ [LINK Table S2](https://docs.google.com/spreadsheets/d/11lF22gUDuUA6rlUR769nCgAX2YykVURy/edit?usp=drive_link&ouid=113479068300354772155&rtpof=true&sd=true)

➔ <u>back to text</u>.

**Supplementary Table S3.** Results of strain GIT, attachment, and invasion assay.

➔ [LINK Table S3](https://docs.google.com/spreadsheets/d/1LpUK2QJ9cFnk62PGjOdaJYd29Z_CzH3F/edit?usp=sharing&ouid=113479068300354772155&rtpof=true&sd=true)

➔ <u>back to text</u>.

**Supplementary Table S4.** Number of reads mapped to the genes of the reference genome and their log-fold change and adjusted p values from the deseq2 analysis.

➔ [LINK Table S4](https://docs.google.com/spreadsheets/d/1nWLhZXi24uRSABJSud3CyMvqKIkXZQ43/edit?usp=sharing&ouid=113479068300354772155&rtpof=true&sd=true)

➔ <u>back to text</u>.

**Supplementary Table S5.** Correspondence of genes and pathways in the selected reference genome; KO = KEGG orthology.

➔ [LINK Table S5](https://docs.google.com/spreadsheets/d/1ndpvw22gglBM-YzghuWLNcT9dfcxzdnC/edit?usp=sharing&ouid=113479068300354772155&rtpof=true&sd=true)

➔ <u>back to text</u>.

**Supplementary Table S6.** Pathways overrepresented in the differential gene expression analysis comparing the low- and high-transforming strains within each of the two induction experiments (SPI1, SPI2); de = genes differentially expressed; geneset = genes assigned to a particular pathway or module.

➔ [LINK Table S6](https://docs.google.com/spreadsheets/d/1Lc01kZlghLTYnTKmrjlaEP97q_-MUmkc/edit?usp=sharing&ouid=113479068300354772155&rtpof=true&sd=true)

➔ <u>back to text</u>.

**Supplementary Table S7.** Median and interquartile range of carbon, nitrogen, phosphorus, and sulfur source utilization by case strains (n=30) and control strains (n=30) and Spearman correlation coefficient (rho) and p-value for the correlation between source utilization and transformation efficiency.

➔ [LINK Table S7](https://docs.google.com/spreadsheets/d/1Omi-yLJn1LS48_sjnIgl4o9E1pWTScAm/edit?usp=sharing&ouid=113479068300354772155&rtpof=true&sd=true)

➔ <u>back to text</u>.

